# Sequence-based Functional Metagenomics Reveals Novel Natural Diversity of Functioning CopA in Environmental Microbiomes

**DOI:** 10.1101/2022.02.12.480192

**Authors:** Wenjun Li, Likun Wang, Xiaofang Li, Xin Zheng, Michael F. Cohen, Yong-Xin Liu

**Author notes:** Corresponding authors (Li X); (Liu Y). Equal contribution: Wenjun Li and Likun Wang contributed equally to this work.

## Abstract

Exploring natural diversity of functional genes/proteins from environmental DNA (eDNA) in a high-throughput fashion remains challenging. In this study, we developed a sequence-based functional metagenomics procedure for mining the diversity of copper resistance gene *copA* in global microbiomes, by combining the metagenomic assembly technology, local BLAST, evolutionary trace analysis (ETA), chemical synthesis, and conventional functional genomics. In total, 88 metagenomes were collected from a public database and subjected to *copA* detection, resulting in 93,899 hits. Manual curation of 1214 hits of high-confidence led to the retrieval of 517 unique CopA candidates, which were further subjected to ETA. Eventually 175 novel *copA* sequences of high-quality were discovered. Phylogenetic analysis showed that almost all of these putative CopA proteins are distantly related to known CopA proteins, with 55 sequences from totally unknown species. Ten novel and three known *copA* genes were chemically synthesized for further functional genomic tests using the Cu-sensitive *Escherichia coli* (Δ*copA*). Growth test and Cu uptake determination showed that five novel clones had positive effects on host Cu resistance and uptake. One recombinant harboring *copA*-like 15 (*copAL15*) successfully restored Cu resistance of host with a substantially enhanced Cu uptake. Two novel *copA* genes were fused with the *gfp* gene and expressed in *E. coli* for microscopic observation. Imaging results showed that they were successfully expressed and their proteins were localized to the membrane. The results here greatly expand the diversity of known CopA proteins, and the sequence-based procedure developed overcomes biases in length, screening methods, and abundance of conventional functional metagenomics.

## Introduction

Knowledge on protein natural diversity is important for both evolutionary and bioengineering studies. Natural diversity of genes/proteins like the DNA-directed RNA polymerase subunit beta (RpoB) [1] and the nitrogenase iron protein (NifH) [2] are widely utilized in microbial phylogenetics, particularly for identifying and describing the nonculturable ‘dark matter’ [3]. The known present-day functional proteins represent a small fraction of the proteins that have arisen over the millions or billions of years of natural selection [4]. High-throughput recovery of the natural diversity of functioning protein variants may pave a way to the quest of how the existing natural proteins differ from random sequences [5], and to protein engineering based on the large-scale library of sequence variants of natural selection instead of directed mutagenesis. Sequence-based enzyme redesign has been shown to be successful in the discovery of esterase and endopeptidase of enhanced activity [6].

For functional genes other than phylo-marker genes, the detection of homologous genes/proteins traditionally relies on the genomics exploration of a pure culture. Expansion of full-genome sequencing greatly enhances our ability to assess the natural diversity of a functional gene/protein, while for some non-ubiquitous cellular functions like metal-resistance, probing their natural diversity remains difficult [7]. Metagenomes contain the full genetic information of eDNA and provide an ideal approach to explore the natural diversity of functional genes/proteins. Pathways to mining functional genes from the metagenomes include the sequence-based and function-based [8]. Function-based screening led to the discovery of novel antibiotic resistance genes [9, 10], biosurfactants [11] and a variety of biocatalysts [12, 13]. Sequence-based functional metagenomics bypasses the limitations of the function-based approaches in the availability of screening methods and redundant isolation [14]. Unfortunately, for many genes, particularly those of large sizes, such as metal transporters, it remains challenging to recover full-length genes from eDNA in a high-throughput manner due to difficulties in PCR detection, degenerated primer design or the availability of known homologs [15].

The core gene for microbial copper resistance, *copA*, is such a gene of large size. It normally possesses low abundance in natural eDNA, and has a limited number of characterized variants to date. Reports on Cu resistance genetic determinants can be traced back to decades ago. Tetaz and Luke reported that plasmid pRJ1104 carried by *Escherichia coli* K-12 conferred enhances Cu resistance [16]. The Cu resistant operon *cop* was first found in *Enterococcus hirae*, which contains regulator genes *copY* and *copZ* encoding a repressor and a chaperone, respectively, as well as the structural genes *copA* and *copB* that mediate Cu transport [17].

CopA is one of the most well-known microbial metal transporters [18]. Studies have shown that CopA was able to protect *Streptococcus suis* through Cu efflux [19]. The lack of CopA can make *E. coli* sensitive to Cu and results in accumulation of Cu^+^ in cells [20]. Regularly, each CopA monomer from *E. coli* binds two Cu^+^ and subsequently transfers them to periplasmic Cu chaperone (CusF) coupled to ATP hydrolysis, thus resulting in the transport of Cu^+^ from cytoplasm to periplasm through the CusCBA trimer protein complex [21]. It is worth noting that the role of CopA was reported to be distinct in different bacteria. For instance, in *E. hirae*, CopA was annotated as a Cu importer [22], while the *Bacillus subtilis* CopA was found to be a Cu exporter [23, 24]. Exploration of more functional CopA from the tremendous reservoir of eDNA may not only facilitate the sequence-based protein design of metal transporters, but also expand the functional diversity of known CopA.

Homology-based annotation has predicted a large amount of novel CopA proteins from eDNA of Cu-contaminated environments, which remains to be verified both bioinformatically and experimentally [25–27]. A pipeline of sequence-based functional metagenomics has been developed, and the high-throughput retrieval of metallothionein genes, a family of short genes encoding Cys-rich metal-binding proteins, from a soil microbiome was realized [7]. Similarly, this procedure was applied in the current study to the exploration of CopA, a much longer metal transporter, from eDNA. To achieve this goal, metagenomes from various environmental microbiomes worldwide were collected from a public database, and subjected to the retrieval of full-length *copA* and functional analysis. Evolutionary trace analysis (ETA) was carried out using a cluster of experimentally-tested CopA proteins to generate sequence features for evaluating the CopA candidates. Ten candidate *copA* genes were randomly selected based on phylogenetic analysis and chemically synthesized for subsequent heterologous expression in Cu-sensitive *E. coli* JW-0473-3 (Δ*copA*). Cu uptake by Cu-sensitive *E. coli* harboring the synthesized *copA* genes was determined. Two clones of *copA* were fused with the green fluorescent protein gene (*gfp*) tag and visualized. Overall, the results here demonstrate the power of sequence-based functional metagenomics in mining or even exhausting the natural diversity of a functional gene in microbiomes. The candidate *copA* detected here may have a distinct mechanism for conferring host Cu resistance.

## Results

### Characteristics of ETA and structure of the reported CopA

Phylogenetic analysis (**Figure 1A**) indicated that the 34 putative CopA proteins were mainly from 14 bacterial species, among which 14 closely related CopA proteins were found to be derived from *Staphylococcus aureus* and another 5 were from *Helcobacter pylori* (**Table S1**). All known CopA function in copper efflux, except for the CopA from *Enterococcus hirae*, which was annotated as copper-importing P-type ATPase. In terms of protein length, CopA generally contain about 800 amino acid residues with the longest one (961 amino acid) was from *Yersinia pestis* **(Figure 1B**). Numbers of heavy metal transporting ATPase (HMA) domain of the 14 groups of CopA ranged from 1 to 3, except for CopA of *Legionella pneumophila* subsp*. pneumophila* with no HMA domain found (**Figure 1B**). Most of the CopA from the 14 bacteria had 2-3 HMA domains, while the CopA from *L. pneumophila* subsp. *pneumophila* had no HMA domains (**Figure 1B**). All the CopA possessed an E1-E2 ATPase domain (**Figure 1B**), which is a Cu binding and efflux structure related to ATP hydrolysis and realizing Cu binding and efflux through conformational variation [28].

**Figure 1.**
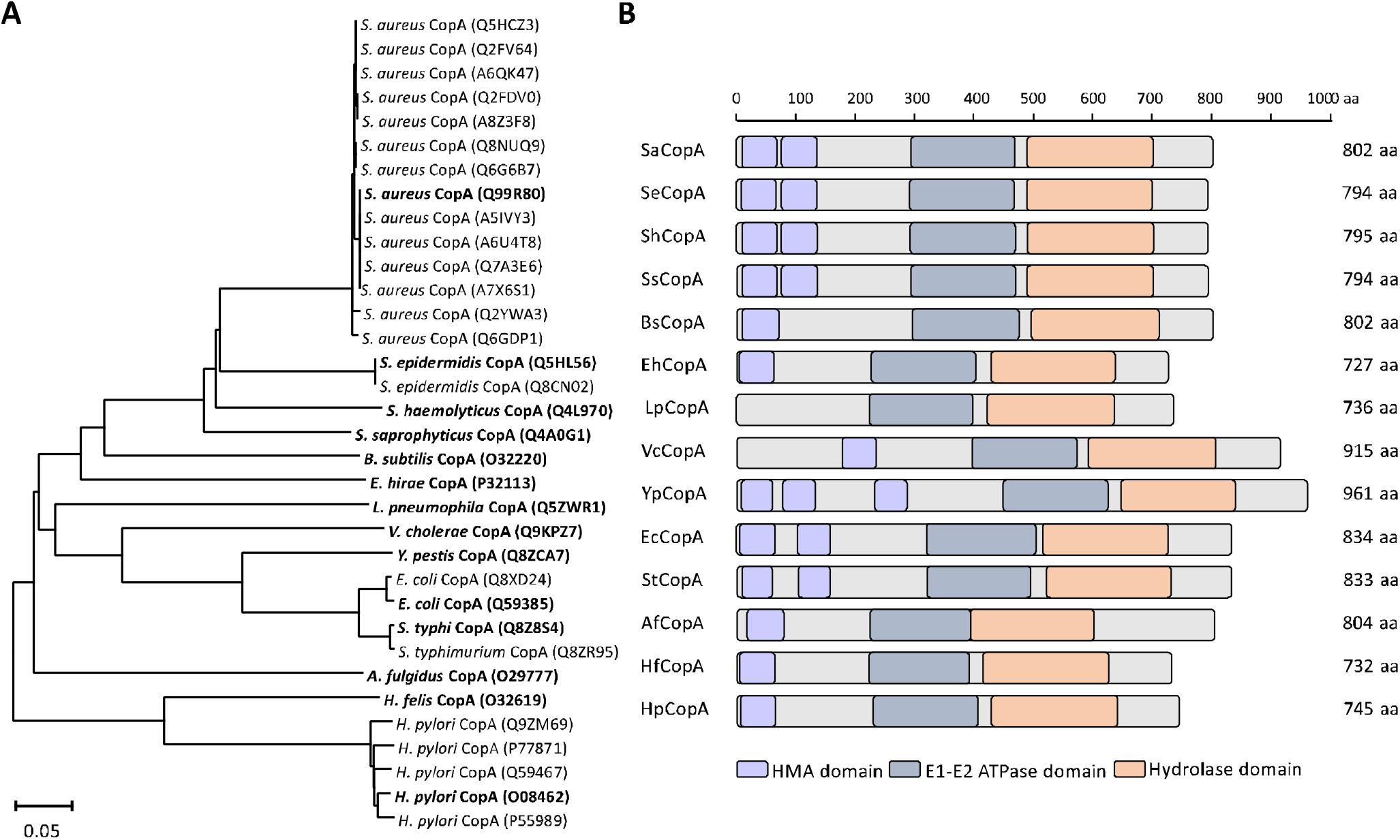
ETA and structural features of reported CopA proteins. **A.** ETA of the 34 known CopA. Amino acid sequences were used to create phylogenetic tree using MEGA 7.0. **B.** Functional domains within the 14 groups of known CopA. Sequences of the 34 known CopA proteins were merged into 14 groups according to species source. Numbers represent the length of the sequences in amino acids. ETA, evolutionary trace analysis. HMA, Heavy Metal transporting ATPase. aa, amino acid.

### Bioinformatics identification of 175 candidate *copA* genes

The assemblage quality varies among the 47 metagenomic datasets, largely due to that the data retrieved from MG-RAST differed in sequencing methods, thus resulting in differences in data size and sequence length (**Table S2**). One low quality dataset mgm4754648 was eliminated from the library and the assembly results of the rest of the 46 metagenomes were included. Eventually 5,500,798 contigs from assemblage and 134,409,173 amino acid sequences from the other 41 datasets were input for local blast.

A total of 93,899 hits were obtained after searching the metagenomic assemblages against the known-CopA database. Then 1,214 returned records of high quality were selected for manual retrieval of CopA candidates from the hits of highest confidence. Through ORF-finder analysis, 517 sequences with the length ranged from 500 to 900 amino acid were preserved and subsequent to transmembrane helices prediction. As a result, predicting by TMHMM and pfam, 315 of them possessed transmembrane helices transmembrane helices. Among the 315 sequences, 222 of them contained metal transport-related ATPase domains (HMA and E1-E2 ATPase). By manual curation of the 222 sequences on their CXXC, HXXH, or CXC amino acid conservative domains, 175 sequences were retrieved.

Taxonomy of the 175 sequences was classified by Kraken 2 (Methods and Table S3). They were found to mainly distribute in five phyla: *Proteobacteria, Actinobacteria, Euryarchaeota, Bacteroidetes* and *Firmicutes* (Figure S1A). Among them 120 sequences belonged to 74 known species, 69 genera, and 47 families (Figure S1A). Other 55 sequences were annotated as unknown species (Table S3). Compare with the 10 reported genera with *copA* genes, 68 (98.6%) novel genera with putative *copA* genes were reported in this study (Figure S1B). As for the species levels, all the 74 species had no overlapped with the reported *copA* genes (Figure S1C). These novel *copA* genes greatly extended the taxonomic diversity of known *copA* genes.

ETA results of the 175 CopA candidates and the 34 known CopA revealed that the sequences were separated into four main branches, however, the 34 known CopA were only in two of the branches. A large proportion of the CopA candidates were located in different developmental branches from the known CopA (**Figure 2**).

**Figure 2.**
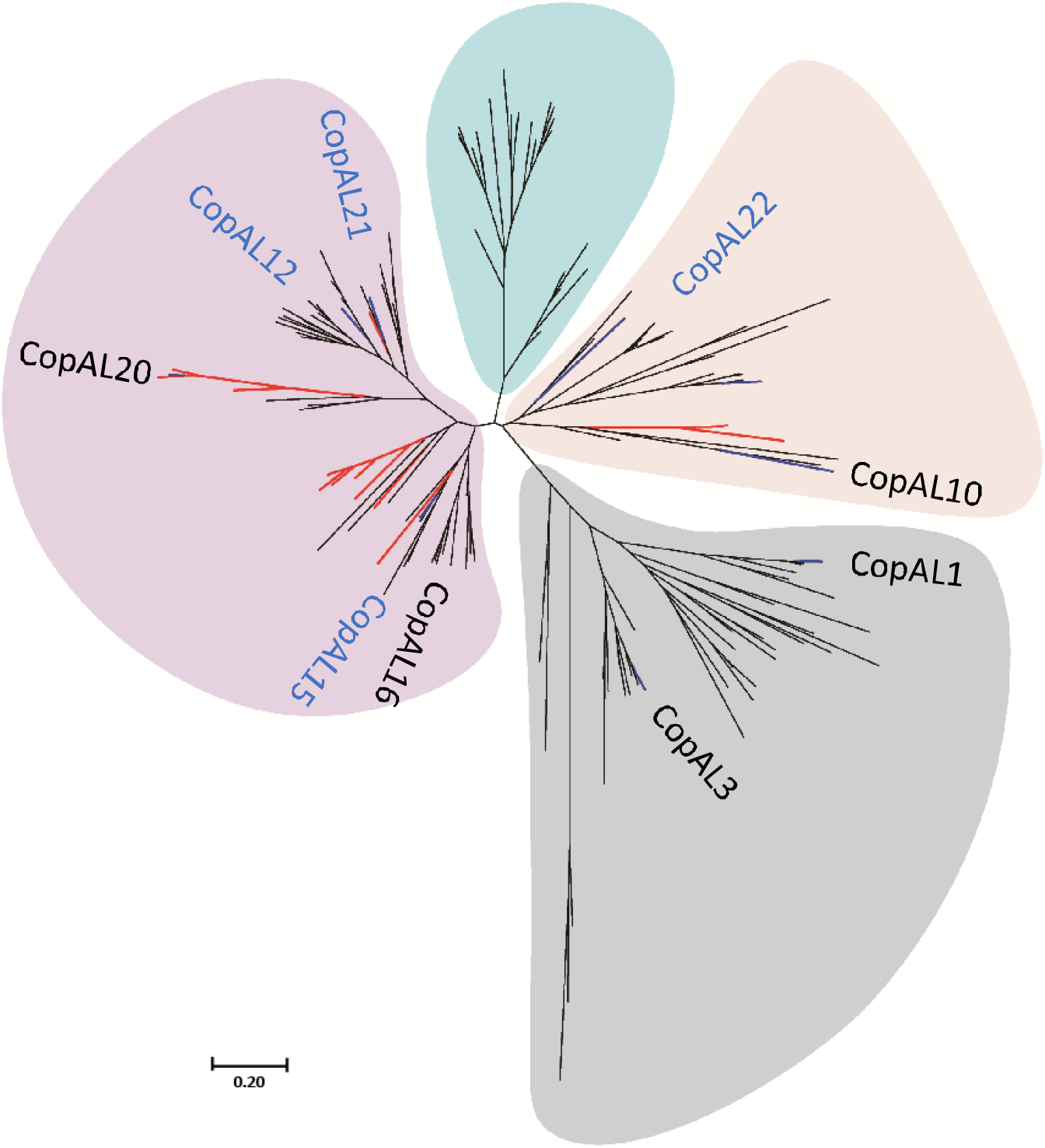
ETA of the 175 candidates and the 34 known CopA proteins. Lines in red are the 34 known CopA, and lines in blue represent the ten genes that were functionally tested via the experiments detailed herein. CopA names in blue are the 5 clones that altered the Cu resistance capacity of the host *Escherichia coli* JW-0473-3 (Δ*copA*) relative to the negative control. The phylogenetic tree of the 175 CopA-like (CopAL) candidates was constructed with MEGA 7.0 using the maximum likelihood method and 1,000 bootstrap replicates. ETA, evolutionary trace analysis.

### Functional genomic verification of 10 selected *copA* genes

Ten *copA*-like genes were selected for chemical synthesis. Amino acid sequences of the 10 selected CopA were back compared with the 34 CopA sequences in the local database using phylogenetic analysis (**Figure 3A**). Overall, Sequences of CopA-like 15 (CopAL15), CopAL20, CopAL10, CopAL16, CopAL12 and CopAL21 were closer to the CopA in local database, while sequences of CopAL22, CopAL6, CopAL1, and CopAL3 were divergent from them and presented independent branches in the phylogenetic tree (**Figure 3A**). More specifically, CopAL6, CopAL1, and CopAL3 were like each other and they were separated from CopA22. In the phylogenetic tree, sequences of CopAL16, CopAL12 and CopAL21 were similar to each other and possessed high homolog with CopA of *Legionella pneumophila* (**Figure 3A**). Additionally, sequences of CopAL15 were more like CopA in *Staphylococcus* spp., *Bacillus subtilis* and *E. hirae*. Notably, the CopA in *E. hirae* was the only one annotated as functioning in Cu import instead of efflux (**Figure 3A**).

**Figure 3.**
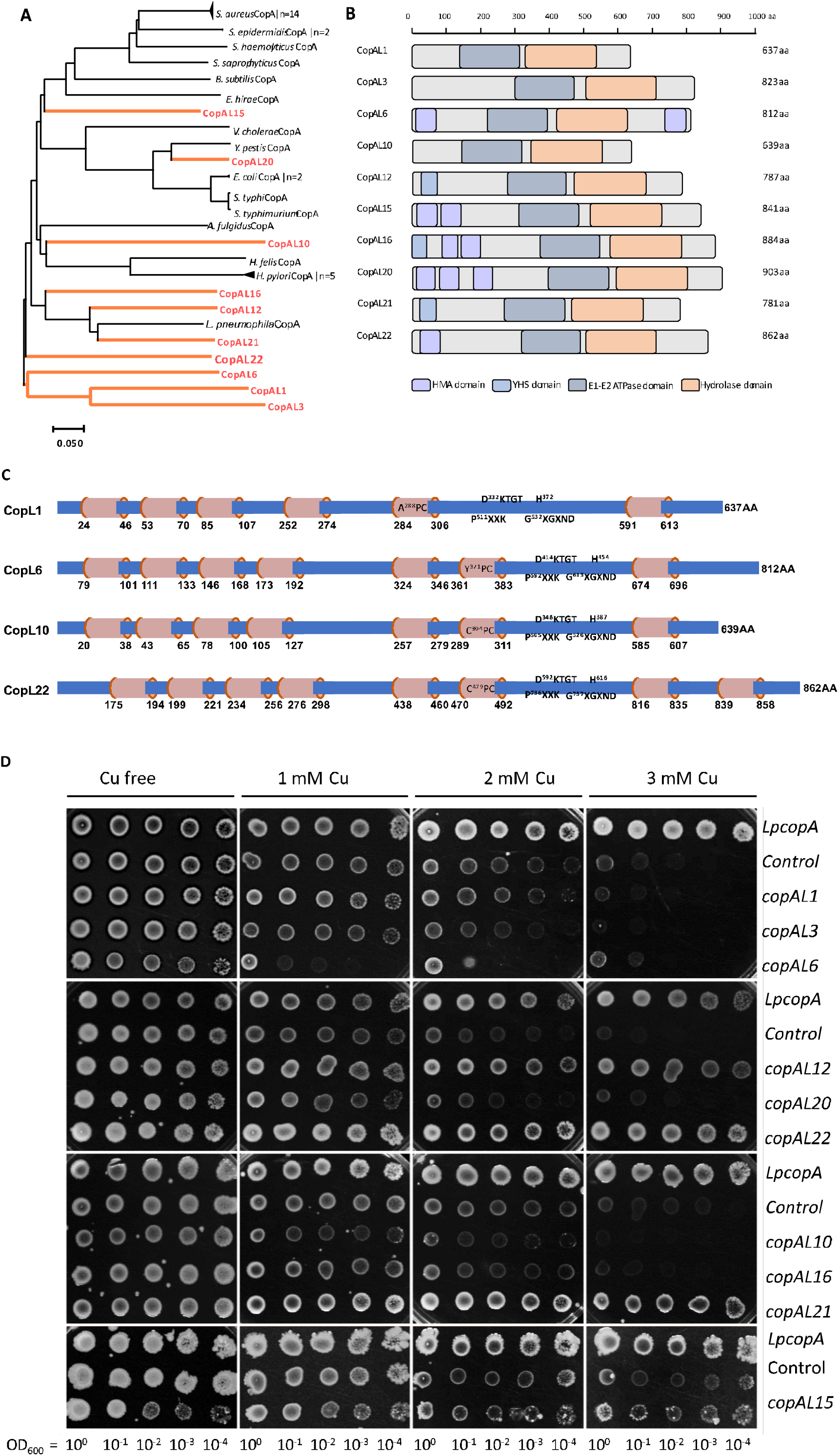
Functional genomic verification of the 10 selected *copA* genes. **A.** Evolutionary trace analysis (ETA) of the 34 putative CopA proteins and the 10 CopA candidates. Amino acid sequences were used to construct a phylogenetic tree using MEGA 7.0. **B.** Functional domains within the 10 selected CopA. **C.** Schematic illustration of the average polypeptide composition of the 10 selected CopA. Among them, CopAL1 represents the composition of both CopAL1 and CopAL3; CopAL22 represents CopAL12, CopAL15, CopAL16, CopAL20, CopAL21, and CopAL22. **D.** Drop assay of the 10 potential CopA proteins. LpCopA: *E. coli* strain JW0473-3 harboring recombinant pTR-Lp*copA*; control: *E. coli* strain JW0473-3 harboring the pTR vector. aa, amino acid.

The length of the 10 putative CopA proteins ranged from 637 to 903 amino acids. All 10 genes contain one E1-E2 ATPase domain and one hydrolase domain, whereas the number of HMA are different among them (**Figure 3B**). The location of the two HMAs in CopAL6 is different from the other CopA, which are found on the two ends of the sequence (**Figure 3B**). Additionally, an YSL (yellow stripe-like) domain that can bind to transition-metal is predicted in CopAL12, CopAL16, and CopAL21 (**Figure 3B**). Notably, the protein sequences of CopAL12 and CopAL16 are highly similar to the known CopA protein LpCopA that lacks an HMA domain.

CopAL12, CopAL15, CopAL16, CopAL20, CopAL21and CopAL22 contain 8 transmembrane domains with a cysteine proline cysteine (CPC) trimer located within the sixth transmembrane domain (**Figure 3C**). CopAL10 has 7 transmembrane domains, and a CPC trimer is also located in the sixth domain (**Figure 3C**). The transmembrane domain prediction of CopAL6 also showed 7 transmembrane domains, while the metal biding site of the sixth transmembrane domain is tyrosine proline cysteine (YPC) trimer. CopAL1 and CopAL3 only possess 6 transmembrane domains, with the fifth transmembrane domain containing an alanine proline cysteine (APC) or a serine proline cysteine (SPC) trimer, respectively (**Figure 3C**). Furthermore, all the 10 synthetic genes possess ATP binding sites, such as monohistidine (H) and aspartate lysine threonine glycine threonine (DKTGT) pentamer.

Ten *copA*-like genes were transformed into *E. coli* JW-0473-3 (Δ*copA*) through a pTR vector. Growth of the negative control was inhibited in 2 mM Cu medium and completely suppressed under 3 mM Cu stress, while growth was not restricted even in 3 mM Cu solid medium for the positive control (**Figure 3D**). Accordingly, function of the 10 candidate genes were classified into three categories, 1) reduced Cu resistance of the host (CopAL6); 2) enhanced Cu resistance of the host (CopAL12, CopAL15, CopAL21 and CopAL22); 3) no change in Cu resistance of the host (CopAL1, CopAL3, CopAL10, CopAL16 and CopAL20). Additionally, Cu-sensitive strains harboring *copAL12, copAL15, copAL21* and *copAL22* showed Cu resistance similar to that of positive control (**Figure 3D**).

### Effects of selected *copA* genes on the host’s growth and metal sorption, and cellular localization of CopA

Growth curves of the *E. coli* JW 0473-3 (Δ*copA*) strains harboring *copAL6, copAL12, copAL15, copAL21, copAL22* along with the positive and negative controls were determined in 2 mM Cu medium (**Figure 4A**). All the five samples reached the stationary stage after 7h incubation. Among them, the growth curves of Cu-sensitive strain harboring *copAL12, copAL15, copAL21*, and *copAL22* were similar with that of the positive control, and all of them grew faster than the negative control, whereas the growth of *E. coli* JW 0473-3 (Δ*copA*) strain harboring *copAL6* was slower than the negative control, indicating expression of *copAL6* inhibited the growth of the sensitive host.

**Figure 4.**
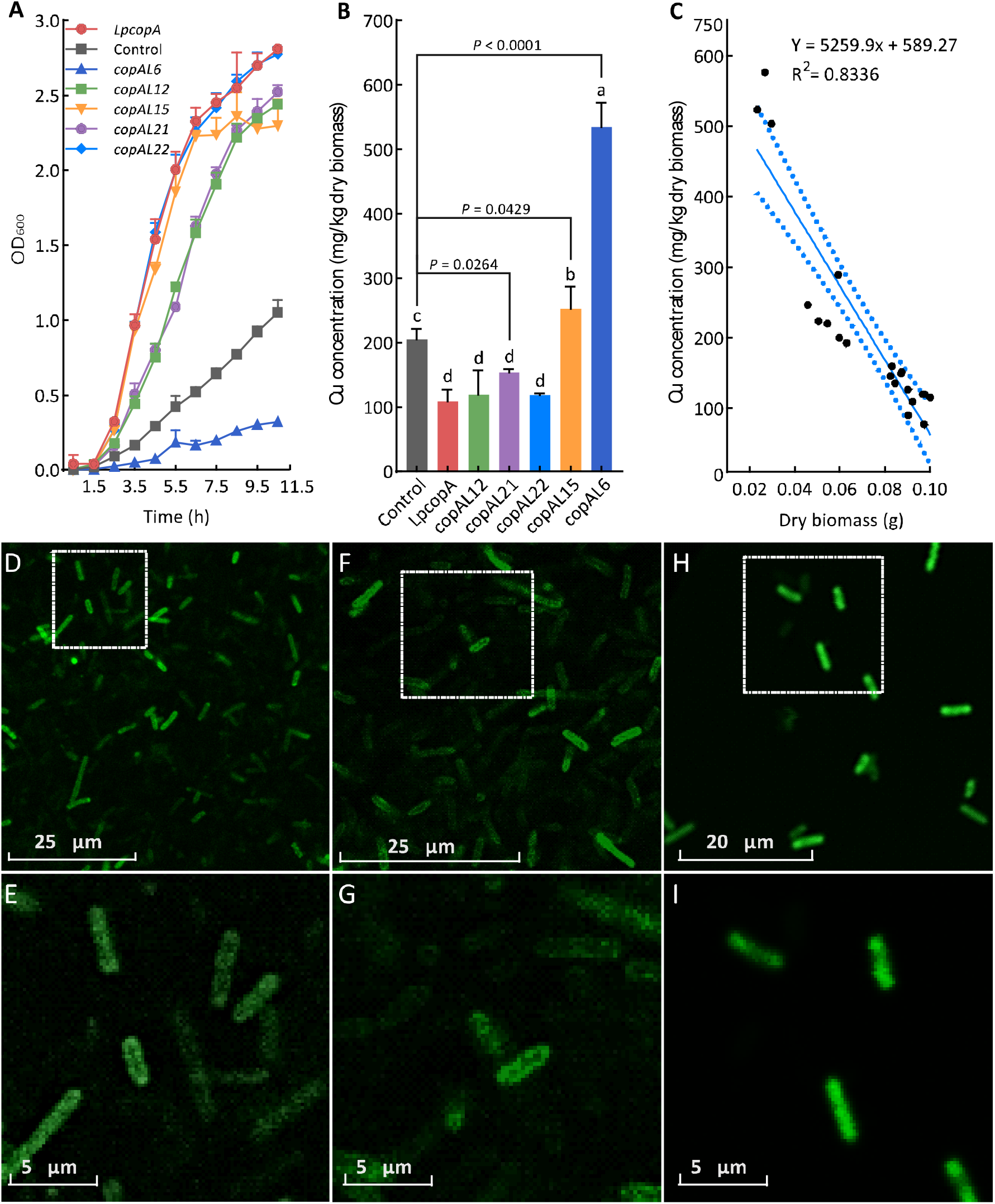
Selected *copA* genes impact the host’s growth and metal sorption, and demonstrate an expected cellular localization. **A.** Growth of strains of *E. coli* JW 0473-3 (Δ *copA*) strains harboring *copAL6, copAL12, copAL15, copAL21, copAL22* along with the positive and negative controls determined in medium with 2mM Cu. **B.** Cu accumulation in *E. coli* JW0473-3 (Δ*copA*) harboring recombinant pTR plasmids, determined by using inductively coupled plasma mass spectrometry (ICP-MS) under 1mM Cu. **C.** Correlation analysis of biomass and Cu accumulation value. LpCopA: *E. coli* JW0473-3 harboring recombinant pTR-Lp*copA*; control: *E. coli* JW0473-3 harboring the pTR vector. **D/E.** Recombinant of *E. coli* strain DH5α harboring a *copAL12-gfp* fusion and green fluorescence was observed located only on the cell membrane. **F**/**G.** Recombinant *E. coli* DH5α strain harboring a *copAL16-gfp* fusion and green fluorescence was observed located only on the cell membrane. **H**/**I.** Recombinant of *E. coli* DH5α harboring *gfp* and green fluorescence was observed throughout the whole cell. Green fluorescence was visualization through confocal laser scanning microscope.

Under 1 mM Cu, sensitive strains harboring *copAL6* and *copAL15* had strong Cu uptake capacities, with bioaccumulation significantly higher comparing to the negative control. Notably, under this Cu stress condition, bioaccumulation of Cu by *E. coli* with *copAL6* was 534. 5 μg/g, which was the highest among all of the *copA*-like genes (**Figure 4B**). In contrast, Cu accumulation by *E. coli* JW 0473-3(Δ*copA*) strain harboring *copAL12, copAL21, copAL22* and the positive control were not significantly different from each other, yet they were all significantly (Fisher’s LSD, *P* ≤ 0.05) lower than the negative control (**Figure 4B**).

A correlation curve was created by plotting the biomass of transformants against Cu accumulation values (**Figure 4C**). A negative correlation was observed between Cu absorption and dry weight with a correlation coefficient (R^2^) of 0.834. Among the ten selected *copA* genes, Cu sensitive strain harboring *copAL6* showed the strongest Cu absorption capacity and a relatively low biomass (**Figure 4C**). Interestingly, both Cu absorption capacity and harvested biomass of *E. coli* harboring *copAL15* were significantly (Fisher’s LSD, *P* ≤ 0.05) higher than the negative control (**Figure 4C**).

Green fluorescence was observed in *E. coli* harboring *copAL12-gfp, copAL16-gfp* and in the positive control (**Figures 4D/F/H**). In addition, strong fluorescence in both recombinants harboring *copAL12-gfp* and *copAL16-gfp* was observed to be localized to the cell membrane, suggesting they may be transmembrane transport proteins (**Figures 4E/G/I**).

## Discussion

In the current study, a sequence-based functional metagenomics procedure was developed to mine the natural diversity of novel CopA proteins from eDNA. The procedure integrated metagenomic mining, ETA, chemical synthesis, and conventional functional genomics using a Cu-sensitive strain (**Figure S1**). The application of this procedure to the exploration of 88 metagenomes from worldwide resulted in the discovery of 175 candidate *copA* genes of high confidence, among which 10 were randomly selected and chemically synthesized for functional genomic tests. Drop assays and growth curve determination showed that 5 clones altered the Cu resistance capacity of the host *E. coli* JW-0473-3 (Δ*copA*) relative to the negative control, among which four (CopAL12, CopAL15, CopAL21 and CopAL22) restored Cu resistance of the sensitive strain and one (CopAL6) reduced the host’s Cu resistance. Interestingly, these five *copA* clones exerted different impacts on Cu accumulation of the host in a manner that was significantly negatively correlated with their dry biomass. Imaging evidence showed that one positive *copA* fused with green fluorescent protein tag was successfully expressed in host cells and probably located in the cell membrane.

CopA belongs to the P_1B-2_ type ATPase family, one of the most well-known metal transport families. Evolutionary analysis of known CopA proteins revealed the conserved metal binding motifs on both terminals [29]. Together with the recent reports on the complete or partial crystal structure of CopA proteins [30–33], this enables the homology-based annotation of novel CopA proteins. It is estimated that CopA abundance in the metagenomes was lower than 0.067%, which is very close to that of natural soil and much lower than Cu-contaminated mine wastes [25]. Annotation of genes in metagenomes has reached a high level of sophistication, while their functional verification, particularly at the high-throughput fashion, is still difficult [34]. In this study, we randomly synthesized and tested 10 full-length *copA* genes from the metagenomes, and to our surprise, 5 of these modified host Cu resistance and uptake of a Cu-sensitive *E. coli* strain. As a transporter is of large-size relative to the other families, the heterologous expression of *copA* is not trivial. The successful detection of 5 candidate functional *copA* genes indicated a high reliability and high possibility of the sequence-based procedure developed here, and we also expect that a large number of functional CopA may present among the 175 candidate CopA proteins. With the lowering of the cost of DNA synthesis, we will gain the ability to apply this method to exhaustively assess the activity of these candidate CopA proteins and also explore loss of function mutations.

Traditionally, screening target genes from eDNA with functional metagenomics approaches involves pressure selection [35]. However, this method was often problematic due to some experimental difficulties, particularly the bias in length of the inserts [36]. Our previous studies explored the possibility of using conventional functional metagenomics to detect novel Cu resistance genes from eDNA, while the results showed that all clones of Cu resistance were not transporter-like genes and with lengths shorter than 1.8 kb [35]. In contrast, metagenomics provides a mean of assessing the total genetic pool of all the microorganisms in a particular environment, which makes it possible to search for large-size functional genes, such as *copA*, without any biases [25, 29]. By means of the new metagenomic pipeline used in the present study, small DNA fragments were assembled into large size contigs which could cover the whole *copA* sequence length of *ca*. 2,400 bp. Again, this study demonstrated the power of sequence-based functional metagenomics in mining large-size functional genes which are difficult for traditional library-based metagenomics.

As mentioned above, although CopA were annotated as Cu efflux proteins in most of the Cu resistant microorganisms, one was found to function in Cu import in *E. hirae* [22]. Traditional gene mining generally involves first obtaining pure cultures of the potential functional microbes, and this may lead to preference for Cu efflux-type CopA proteins. The procedure used in our study does not rely on the screening of Cu resistance in detecting candidate CopA proteins, and thus overcomes the bias for efflux functions. Accordingly, in our results, we found a novel CopA protein with a function of Cu import thereby increasing the host’s Cu sensitivity. Considering that all known Cu resistance systems like Cop, Pco and Cus are of low abundance in natural environments [37, 38], use of traditional metagenomics that relies on PCR cloning and library construction is not a realistic means for probing the natural diversity of these Cu resistance genes.

Although heterologous expression of targeted genes in a host can be extremely challenging, we achieved a relatively high success rate using a domesticated *E. coli* host strain [39]. Growth test and metal uptake determination showed successful detection of five candidate CopA proteins, demonstrating a 50% rate of detecting positive clones from the eDNA. In addition to the physiological evidence, imaging results based on GFP-fusion visualization further confirmed the successful expression of *copAL12* in the host and revealed the likely localization of these proteins to the cell membrane. In contrast, we also found that a candidate CopA (CopAL16) was successfully transcribed and translated in the host based on the GFP-fusion visualization evidence but which did not alter host Cu resistance (**Figures 4F** and **4G**). In some cases, foreign DNA can be expressed in the heterologous host but the gene function be silent due to the lack of chaperones required for proper protein folding [8]. A protein that did not display antimicrobial activity in *E. coli* host did confer this activity to a *Ralston metallidurans* host [40], thus indicating the importance of using additional heterologous hosts to identify active clones that fail to express in the standard *E. coli* host [41]. We thus anticipate that the procedure developed here may be able to probe with a high rate of success the natural diversity of CopA and other proteins involved in metal transport.

## Material and methods

### Environmental metagenomes collection and assembly

Eighty-eight environmental metagenomic datasets were collected from our previous studies as well as the MG-RAST (Metagenomics Rapid Annotation using Subsystem Technology) server (**Figure 5**). The metagenomic datasets represented eDNA from a global diversity of habitats including farmland soil, forest soil, wastewater, contaminated soil, mine drainage, mine tailings, and ocean water (**Figure 5**). Among the 88 datasets, 47 of them were quality-controlled DNA sequences and host DNA was already removed (**Table S2**), thus data were assembled using the assembly module in Metawrap v. 1.2.1 through their in-house scripts [42] (https://github.com/ebg-lab/CopA.git). The other 41 datasets were amino acid sequences, thus allowing for similarity comparison through BLASTP to be directly performed. Total bases counts, total read counts, number of contigs, the largest contig length and N50 were recorded (**Table S2**). Additionally, in order to facilitate the use of this method by colleagues, the detailed step-by-step protocols are in the supplementary protocols “A manual for detecting CopA of high confidence from metagenomes”.

**Figure 5.**
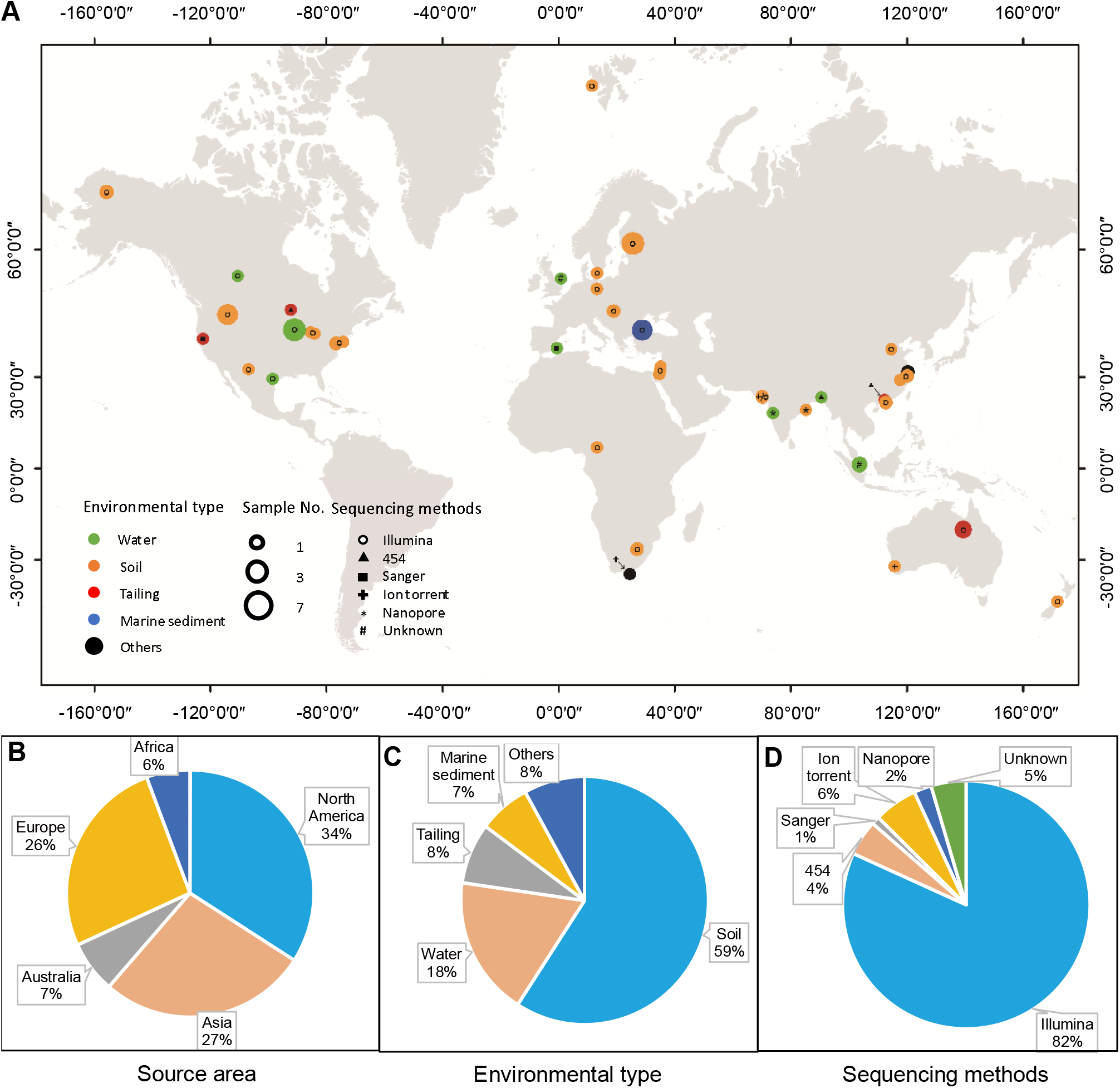
Basic information of the 88 metagenomic datasets. **A.** Geographic distribution, the type of environmental source and sequencing method of the 88 datasets. The color of points refers to the type of environmental sources of the datasets. Circle size is used to distinguish number of datasets in each location. Sequencing method is represented by the graphical shape in the points. Proportion of datasets among continents (**B)**, environmental types (**C)**, and sequencing methods (**D)**.

### Known CopA database construction for local BLAST

Thirty-four amino acid sequences marked with ‘manually annotated’ were retrieved from the Uniprot database (https://www.uniprot.org/) with ‘CopA’ as an entry. These sequences were experimentally characterized for either protein structure or metal-resistance function. The 34 sequences were mainly from 14 microorganisms, which were listed in **Table** S**1**. Data were re-formatted using makeblastdb from blast v2.2.31 (paramters ‘-in nucleotide.fa -dbtype nucl’ for nucleotide, and “-in protein.fa -dbtype prot” for amino acid) to create index of database. The phylogenetic relationship of the 34 CopA sequences was constructed with MEGA 7.0 [43] using the maximum likelihood method and 1,000 bootstrap replicates. Multiple sequence alignment was performed using ClustalW [44], and p-distance was calculated. All positions with less than 50% site coverage (namely ≥ 50% alignment gaps, missing data, or fuzzy bases) were eliminated. Domain and motif analysis of the 34 CopA sequences were followed by the same procedure as for the candidate CopA below.

### Local BLAST for candidate *copA* detection

The environmental metagenomes were searched against the CopA database via local BLAST [45], which was done on a Linux system computer equipped with dual-core 2.2 × 2 GHz CPU and 192 G RAM. Briefly, nucleotide sequences with length greater than 2000 bp in the 47 assembled metagenomic datasets and amino acid sequences with length greater than 700 aa in the other 41 datasets were aligned against the CopA local database using the BLASTX and BLASTP method, respectively. Only those matches having an E-value of ≤ 1.0 × 10^-6^ were recorded.

The matches obtained from blast were subjected to search for open reading frames (ORFs) using ORF finder (https://www.ncbi.nlm.nih.gov/orffinder/) with ATG as the initiation codon. Candidate ORFs with the lengths ranging from 500 to 900 amino acids were selected and subjected to the prediction of transmembrane helices using the TMHMM (http://www.cbs.dtu.dk/services/TMHMM-2.0/) online analysis platform. Functional domains were then predicted using Pfam (http://pfam.xfam.org/) with an E-value of ≤ 1e-6, and sequences without metal transporting domains were eliminated. Eventually, *copA*-like sequences of high confidence with CXXC, HXXH or CXC conserved metal-binding domains were manually retrieved. A phylogenetic tree showing the evolutional relationship of the candidate CopA proteins was constructed using the abovementioned method.

### Bacterial strains and cultural conditions for functional genomics

A pTR vector carrying endonuclease sites *Pst* I and *Kpn* I was used as an expression vector in this study (**Figure** S**2**) [7], and Cu-sensitive *E.coli* JW0473-3 (Δ*copA*) was used as the host. *E. coli* JW0473-3 harboring the pTR vector was set as the negative control, and the one harboring recombinant pTR-Lp*copA* was used as a positive control. Common *E. coli* strain DH5α (F-f801acZDM15 D (*lacZYA-argF*) U169 recA1 endA1 hsdR17 (rk-, mk+) phoA supE44 thi-1 gyrA96 relA1 l-, Invitrogen, USA) was used to store the recombinant plasmids and for heterologous expression of green fluorescent protein-fused genes and imaging. Green fluorescence was observed through a confocal laser scanning microscope (Leica TCS-SP8, Leica Microsystems).

Luria-Bertani (LB) medium supplied with 100 mg mL^-1^ ampicillin and 50 mg mL^-1^ kanamycin was used to select *E. coli* JW0473-3 (Δ*copA*) recombinants.

### Taxonomy classification, comparison and visualization

Taxonomic classification of the novel *copA* genes was performed using Kraken 2.1.1 [46] based on the NCBI taxonomy database version 20210120. Then the NCBI taxonomy IDs were converted into standard 7 ranks table by Taxonkit 0.7.0 [47]. The formatted taxonomic information is in Table S3. The data process for Cladograms were follows the guide of EasyAmplicon 1.14 [48], and finally visualized by ImageGP webserver 1.0 [49] by calling GraPhlAn v.0.9.7 [50]. The comparison known and novel taxonomy of *copA* genes is analyzed in Evenn webserver [51]

### Sequences synthesis

According to the phylogenetic analysis result of the 34 known CopA and 175 CopA candidates, ten sequences with a length of 2,000-2,700 bp were randomly selected from the candidate *copA* genes and subjected to artificial DNA synthesis as well as subsequent functional verification. Codon preference of the host, secondary structure of mRNA and GC content were considered [52] for DNA synthesis to improve the expression efficiency in the host *E. coli*. The recombinant pTR-*copA* were re-digested with *Pst*I and *Kpn*I restriction enzymes to examine the successfulness of insertion and *copA-like* sequences were double checked by sequencing on Sanger platform. pTR vector recombined with *copA*-like sequences and *gfp* were transformed into common *E. coli* DH5α for visualization of green fluorescence.

A phylogenetic tree of the 10 *copA-like* candidates and 34 reported *copA* sequences was created using MEGA 7.0 with the maximum likelihood method and 1,000 bootstrap replicates.

### Drop assay for Cu resistance screening

The function of the 10 *copA*-like genes were verified via drop assay experiments described in our previous study [7]. In brief, recombinant pTR vector harboring *copA*-like genes were first transformed into *E. coli* JW0473-3 (Δ*copA*) by using a CaCl_2_-based chemical transformation method. The transformed cells were then incubated at 37 °C in liquid LB medium supplied with 100 mg mL^-1^ ampicillin and 50 mg mL^-1^ kanamycin overnight. Cells were harvested by centrifugation, and then re-suspended in water to obtain an OD_600_ of 1.0. A gradient dilution down to 10^-4^ was performed. A total of 3 μl dilution was inoculated onto LB plates containing different concentrations of Cu (1, 2, 3 and 4 mM Cu^2+^, as CuSO_4_·5H_2_O). MIC (minimum inhibitory concentration) of the sensitive strains harboring recombinant pTR was determined.

### Growth test

Cu-sensitive strains harboring recombinant pTR were incubated in liquid LB medium overnight and adjusted the initial OD_600_ to 0.1. Growth curve of the recombinants and controls were measured by incubating in liquid LB medium with 2 mM Cu at 37 °C. The concentration of each culture was measured by a biophotometer (Eppendorf, Germany) at a 30-minute interval for 8 h.

### Biomass and metal sorption determination

Cu-sensitive strains harboring the recombinant pTR plasmids were harvested for metal content determination by using inductively coupled plasma mass spectrometry (ICP-MS) (Thermo Fisher Scientific, USA). *E. coli* were incubated in liquid LB medium overnight and adjusted the initial OD_600_ value to 0.02. After another 8 h of incubation in liquid LB medium with 1mM Cu at 37 °C, cells were collected by centrifugation at 4000×g. Cells pellets were rinsed by ultra-pure water and subsequently dried, weighed, and digested by using 8 mL of 65% HNO_3_ and 2 mL 70% HClO_4_. The digested mixture was dissolved in 50 mL Millipore-filtered water, and then the metal content was measured using ICP-OES (Optima 7000 DV, Perkin Elmer, MA, USA). Certified reference material laver (GWB10023 Certified by the Institute of Geophysical and Geochemical Exploration, China) was used as a standard reference material for determining of Cu concentrations.

### Statistical analysis

All comparisons were subjected to analysis of variance (ANOVA) using SAS (version 9.4; SAS Institute, Cary, NC) GLM model.

Means separation was conducted by using the Fisher’s LSD (Least Significant Difference) test with *P* ≤ 0.05 considered significant.

## Acknowledgments

XL was supported by the National Natural Science Foundation of China (No. 41877414), the National Key R&D Program of China (No.2018YFD0800306), and the Hebei Provincial Science Fund for Distinguished Young Scholars (No. D2018503005). XZ was supported by the National Natural Science Foundation of China (No. 31700228). YX was supported by the National Natural Science Foundation of China (U21A20182) and the Youth Innovation Promotion Association CAS (2021092).

## CRediT author statement

Wenjun Li: Validation, Formal analysis, Writing - original draft, Visualization. Xiaofang Li: Conceptualization, Writing - original draft, Visualization, Writing - review & editing. Yong-Xin Liu: Writing - review & editing, Analysis, Visualization. Likun Wang: Formal analysis, Writing – original draft, Writing - review & editing, Visualization. Xin Zheng: Experiment, Methodology, Writing - original draft, Writing - review & editing, Visualization. Michael F. Cohen: Writing - review & editing. All authors read and approved the final manuscript.

## Supplementary material

**Figure S1** Taxonomy of novel *copA* genes. **A.** Taxonomy of novel *copA* genes in Cladogram. The taxonomy of *copA* sequences were classified by Kraken2 based on NCBI database. The colored background represents Phylum. The labels are Family. Only the sequences have known family were showed in this figure. B. Overlapping genus of known and novel *copA*. C. Overlapping species of known and novel *copA*. The taxonomy of 175 novel *copA* gene are listed in Table S3.

**Figure S2** Experimental procedures for mining the natural diversity of CopA.

**Figure S3** A graphic map of *Escherichia coli* expression vector pUC19-Prrn-T7g10-rrnBT1 (PTR).

**Table S1** Basic information of reported CopA from the fourteen bacterial species.

**Table S2** Assemblage and blast results of the 88 metagenomic datasets.

**Table S3** Taxonomy of 175 novel copA genes by Kraken2 classification.

**Supplementary protocols** A manual for detecting CopA of high confidence from metagenomes.

## REFERENCES

[1] Wang Z, Wu M: A phylum-level bacterial phylogenetic marker database. Mol Biol Evol 2013, 30(6):1258–1262.

[2] Kapili BJ, Dekas AE: PPIT: an R package for inferring microbial taxonomy from nifH sequences. Bioinformatics 2021, 37(16):2289–2298

[3] Wu D, Hugenholtz P, Mavromatis K, Pukall R, Dalin E, Ivanova NN, et al.: A phylogeny-driven genomic encyclopaedia of Bacteria and Archaea. Nature 2009, 462(7276):1056–1060.

[4] Cole MF, Gaucher EA: Utilizing natural diversity to evolve protein function: applications towards thermostability. Curr Opin Chem Biol 2011, 15(3):399–406.

[5] Yu JF, Cao ZX, Yang YD, Wang CL, Su ZD, Zhao YW, et al. Natural protein sequences are more intrinsically disordered than random sequences. Cell Mol Life Sci 2016, 73(15):2949–2957.

[6] Lutz S. Beyond directed evolution-semi-rational protein engineering and design. Curr Opin Biotech 2010, 21(6):734–743.

[7] Li X, Islam MM, Chen L, Wang L, Zheng X. Metagenomics-guided discovery of potential bacterial metallothionein genes from the soil microbiome that confer Cu and/or Cd resistance. Appl Environ Microbiol 2020, 86(9).

[8] Uchiyama T, Miyazaki K. Functional metagenomics for enzyme discovery: challenges to efficient screening. Curr Opin Biotech 2009, 20(6):616–622.

[9] Berglund F, Osterlund T, Boulund F, Marathe NP, Larsson DGJ, Kristiansson. Identification and reconstruction of novel antibiotic resistance genes from metagenomes. Microbiome 2019, 7(1):52

[10] dos Santos DFK, Istvan P, Quirino BF, Kruger RH. Functional metagenomics as a tool for identification of new antibiotic resistance genes from natural environments. Microb Ecol 2017, 73(2):479–491.

[11] Williams W, Trindade M. Metagenomics for the discovery of novel biosurfactants. In: Functional metagenomics: Tools and applications. Edited by Charles TC, Liles MR, Sessitsch A. Cham: Springer International Publishing; 2017: 95–117.

[12] Tiwari R, Nain L, Labrou NE, Shukla P. Bioprospecting of functional cellulases from metagenome for second generation biofuel production: a review. Crit Rev Microbiol 2018, 44(2):244–257.

[13] Armstrong Z, Mewis K, Liu F, Morgan-Lang C, Scofield M, Durno E, et al. Metagenomics reveals functional synergy and novel polysaccharide utilization loci in the Castor canadensis fecal microbiome. ISME J 2018, 12(11):2757–2769.

[14] Jeffries J, Dawson N, Orengo C, Moody TS, Ward JM. Metagenome mining: a sequence directed strategy for the retrieval of enzymes for biocatalysis. ChemistrySelect 2016, 1(10):2217–2220.

[15] Das S, Sen M, Saha C, Chakraborty D, Das A, Banerjee M, Seal A. Isolation and expression analysis of partial sequences of heavy metal transporters from Brassica juncea by coupling high throughput cloning with a molecular fingerprinting technique. Planta 2011, 234(1):139–156.

[16] Tetaz TJ, Luke RK. Plasmid-controlled resistance to copper in *Escherichia coli*. J Bacteriol 1983, 154(3):1263–1268.

[17] Wunderli-Ye H, Solioz M. Copper homeostasis in *Enterococcus hirae*. Adv Exp Med Biol 1999, 448:255–264.

[18] Aguila-Clares B, Castiblanco LF, Quesada JM, Penyalver R, Carbonell J, Lopez MM, et al. Transcriptional response of *Erwinia amylovora* to copper shock: in vivo role of the copA gene. Mol Plant Pathol 2018, 19(1):169–179.

[19] Zheng C, Jia M, Lu T, Gao M, Li L. CopA protects *Streptococcus suis* against copper toxicity. Int J Mol Sci 2019, 20(12).

[20] Petersen C, Moller LB. Control of copper homeostasis in *Escherichia coli* by a P-type ATPase, CopA, and a MerR-like transcriptional activator, CopR. Gene 2000, 261(2):289–298.

[21] Padilla-Benavides T, George Thompson AM, McEvoy MM, Arguello JM. Mechanism of ATPase-mediated Cu+ export and delivery to periplasmic chaperones: the interaction of *Escherichia coli* CopA and CusF. J Biol Chem 2014, 289(30):20492–20501.

[22] Lu ZH, Dameron CT, Solioz M. The *Enterococcus hirae* paradigm of copper homeostasis: copper chaperone turnover, interactions, and transactions. Biometals 2003, 16(1): 137–143.

[23] Radford DS, Kihlken MA, Borrelly GP, Harwood CR, Le Brun NE, Cavet JS. CopZ from *Bacillus subtilis* interacts in vivo with a copper exporting CPx-type ATPase CopA. FEMS Microbiol Lett 2003, 220(1):105–112.

[24] Steunou AS, Durand A, Bourbon ML, Babot M, Tambosi R, Liotenberg S, Ouchane S. Cadmium and copper cross-tolerance. Cu(+)alleviates Cd(2+)toxicity, and Both cations target heme and chlorophyll biosynthesis pathway in *Rubrivivax gelatinosus*. Front Microbiol 2020, 11:893

[25] Li X, Zhu YG, Shaban B, Bruxner TJ, Bond PL, Huang L. Assessing the genetic diversity of Cu resistance in mine tailings through high-throughput recovery of full-length copA genes. Sci Rep 2015, 5:13258.

[26] Liu JL, Yao J, Zhu X, Zhou DL, Duran R, Mihucz VG, et al. Metagenomic exploration of multi-resistance genes linked to microbial attributes in active nonferrous metal(loid) tailings. Environ Pollut 2020, 273:115667.

[27] Martin C, Stebbins B, Ajmani A, Comendul A, Hamner S, Hasan NA, et al. Nanopore-based metagenomics analysis reveals prevalence of mobile antibiotic and heavy metal resistome in wastewater. Ecotoxicology 2021, 30(8), 1572–1585

[28] Andersson M, Mattle D, Sitsel O, Klymchuk T, Nielsen AM, Moller LB, et al. Copper-transporting P-type ATPases use a unique ion-release pathway. Nat Struct Mol Biol 2014, 21(1):43-+.

[29] Smith AT, Smith KP, Rosenzweig AC. Diversity of the metal-transporting P-1B-type ATPases. J Biol Inorg Chem 2014, 19(6):947–960.

[30] Gonzalez-Guerrero M, Arguello JM. Mechanism of Cu+-transporting ATPases: soluble Cu+ chaperones directly transfer Cu+ to transmembrane transport sites. Proc Natl Acad Sci U S A 2008, 105(16):5992–5997.

[31] Lubben M, Portmann R, Kock G, Stoll R, Young MM, Solioz M. Structural model of the CopA copper ATPase of *Enterococcus hirae* based on chemical cross-linking. Biometals 2009, 22(2):363–375.

[32] Rensing C, Fan B, Sharma R, Mitra B, Rosen BP. CopA: An *Escherichia coli* Cu(I)-translocating P-type ATPase. Proc Natl Acad Sci U S A 2000, 97(2):652–656.

[33] Gourdon P, Liu XY, Skjorringe T, Morth JP, Moller LB, Pedersen BP, et al. Crystal structure of a copper-transporting P_IB_-type ATPase. Nature 2011, 475(7354):59–64.

[34] Hess M, Sczyrba A, Egan R, Kim TW, Chokhawala H, Schroth G, et al. Metagenomic discovery of biomass-degrading genes and genomes from Cow Rumen. Science 2011, 331(6016):463–467.

[35] Xing C, Chen J, Zheng X, Chen L, Chen M, Wang L, et al.: Functional metagenomic exploration identifies novel prokaryotic copper resistance genes from the soil microbiome. Metallomics 2020, 12(3):387–395.

[36] Crameri R, Suter M. Display of biologically-active proteins on the surface of filamentous phages - a cDNA cloning system for selection of functional gene-products linked to the genetic Information responsible for their production. Gene 1993, 137(1):69–75.

[37] Outten FW, Huffman DL, Hale JA, O’Halloran TV. The independent *cue* and *cus* systems confer copper tolerance during aerobic and anaerobic growth in *Escherichia coli*. J Biol Chem 2001, 276(33):30670–30677.

[38] Li LG, Cai L, Zhang XX, Zhang T. Potentially novel copper resistance genes in copper-enriched activated sludge revealed by metagenomic analysis. Appl Microbiol Biotechnol 2014, 98(24):10255–10266.

[39] Banik JJ, Brady SF. Recent application of metagenomic approaches toward the discovery of antimicrobials and other bioactive small molecules. Curr Opin Microbiol 2010, 13(5):603–609.

[40] Craig JW, Chang FY, Brady SF. Natural products from environmental DNA hosted in *Ralstonia metallidurans*. ACS Chem Biol 2009, 4(1):23–28.

[41] Craig JW, Chang FY, Kim JH, Obiajulu SC, Brady SF. Expanding small-molecule functional metagenomics through parallel screening of broad-host-range cosmid environmental DNA libraries in diverse Proteobacteria. Applied and Environmental Microbiology 2010, 76(5):1633–1641.

[42] Uritskiy GV, DiRuggiero J, Taylor J. MetaWRAP—a flexible pipeline for genome-resolved metagenomic data analysis. Microbiome 2018, 6(1):158.

[43] Kumar S, Stecher G, Tamura K. MEGA7: molecular evolutionary genetics analysis version 7.0 for bigger datasets. Mol Biol Evol 2016, 33(7):1870–1874.

[44] Thompson JD, Higgins DG, Gibson TJ. Clustal-W - improving the sensitivity of progressive multiple sequence alignment through sequence weighting, position-specific gap penalties and weight matrix Choice. Nucleic Acids Research 1994, 22(22):4673–4680.

[45] Camacho C, Coulouris G, Avagyan V, Ma N, Papadopoulos J, Bealer K, et al. BLAST+: architecture and applications. BMC Bioinformatics 2009, 10:421.

[46]. Wood DE, Lu J, Langmead B: Improved metagenomic analysis with Kraken 2. Genome Biology 2019, 20(1):257.

[47]. Shen W, Ren H: TaxonKit: A practical and efficient NCBI taxonomy toolkit. Journal of Genetics and Genomics 2021, 48(9):844–850.

[48]. Liu Y-X, Qin Y, Chen T, Lu M, Qian X, Guo X, Bai Y: A practical guide to amplicon and metagenomic analysis of microbiome data. Protein & Cell 2021, 12(5):315–330.

[49]. Chen T, Liu Y-X, Huang L: ImageGP: An easy-to-use data visualization web server for scientific researchers. iMeta 2022, 1(1):e5.

[50]. Asnicar F, Weingart G, Tickle TL, Huttenhower C, Segata N: Compact graphical representation of phylogenetic data and metadata with GraPhlAn. PeerJ 2015, 3:e1029.

[51]. Chen T, Zhang H, Liu Y, Liu Y-X, Huang L: EVenn: Easy to create repeatable and editable Venn diagrams and Venn networks online. Journal of Genetics and Genomics 2021, 48(9):863–866.

[52] McPherson DT. Codon preference reflects mistranslational constraints: a proposal. Nucleic Acids Res 1988, 16(9):4111–4120.

