## Supplementary material for "Sequence-based Functional Metagenomics Reveals Novel Natural Diversity of Functioning CopA in Environmental Microbiomes": Figure S

*Running Head: Li W et al / Natural diversity of copper resistance genes from eDNA*

* Corresponding authors

 (Li X); (Liu Y)

#Equal contribution

Wenjun Li and Likun Wang contributed equally to this work. Author order was determined based on contributions.

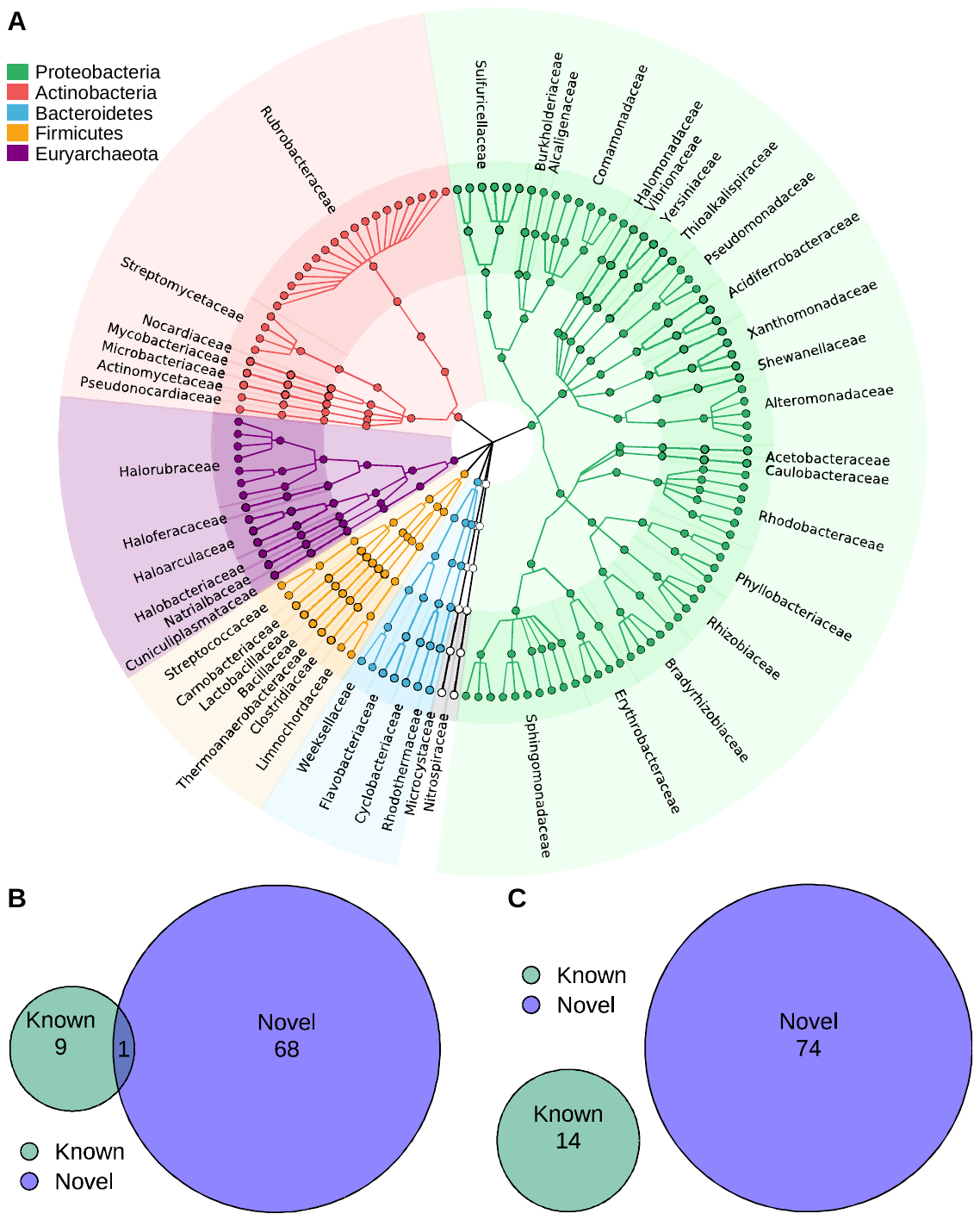

**Figure S1 Taxonomy of novel *copA* genes.**

**A.** Taxonomy of novel *copA* genes in Cladogram. The taxonomy of *copA* sequences were classified by Kraken2 based on NCBI database. The colored background represents Phylum. The labels are Family. Only the sequences have known family were showed in this figure. B. Overlapping genus of known and novel *copA*. C. Overlapping species of known and novel *copA*. The taxonomy of 175 novel *copA* gene are listed in Table S1.

**Figure S2 Experimental procedures for mining the natural diversity of CopA.**

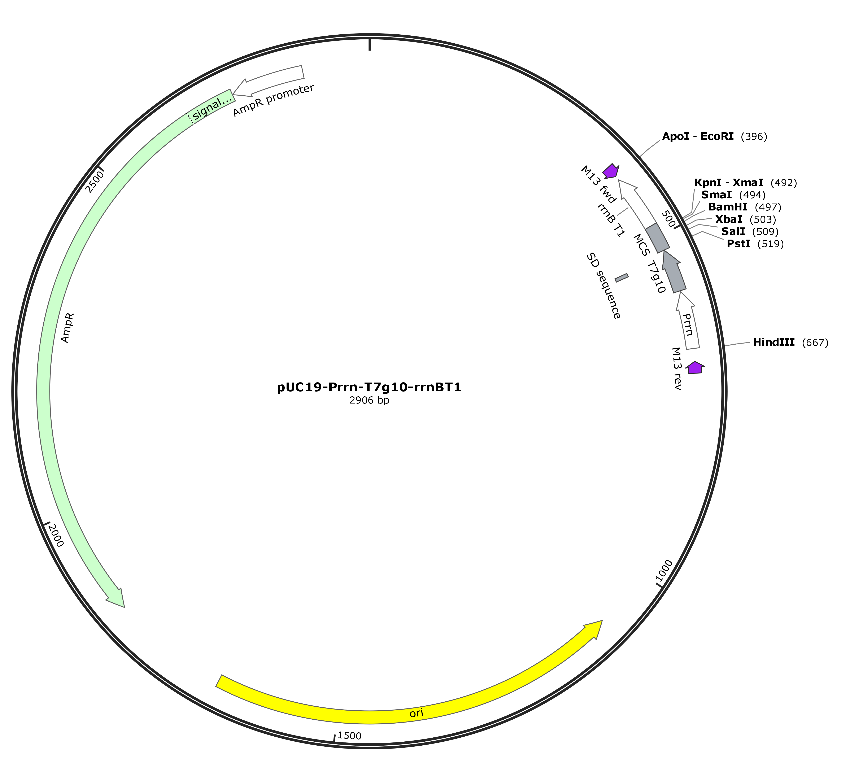

**Figure S3 Map of *Escherichia coli* expression vector pUC19-Prrn-T7g10-rrnBT1 (PTR).**

**REFERENCE**

1.	Odermatt A, Suter H, Krapf R, Solioz M: **Primary structure of two P-type ATPases involved in copper homeostasis in Enterococcus hirae**. *J Biol Chem* 1993, **268**(17):12775-12779.

2.	Mandal AK, Cheung WD, Arguello JM: **Characterization of a thermophilic P-type Ag+/Cu+-ATPase from the extremophile Archaeoglobus fulgidus**. *J Biol Chem* 2002, **277**(9):7201-7208.

3.	Padilla-Benavides T, George Thompson AM, McEvoy MM, Arguello JM: **Mechanism of ATPase-mediated Cu+ export and delivery to periplasmic chaperones: the interaction of Escherichia coli CopA and CusF**. *J Biol Chem* 2014, **289**(30):20492-20501.

4.	Rensing C, Fan B, Sharma R, Mitra B, Rosen BP: **CopA: An Escherichia coli Cu(I)-translocating P-type ATPase**. *Proc Natl Acad Sci U S A* 2000, **97**(2):652-656.

5.	Sitthisak S, Knutsson L, Webb JW, Jayaswal RK: **Molecular characterization of the copper transport system in Staphylococcus aureus**. *Microbiology (Reading)* 2007, **153**(Pt 12):4274-4283.

6.	Radford DS, Kihlken MA, Borrelly GP, Harwood CR, Le Brun NE, Cavet JS: **CopZ from Bacillus subtilis interacts in vivo with a copper exporting CPx-type ATPase CopA**. *FEMS Microbiol Lett* 2003, **220**(1):105-112.

7.	Espariz M, Checa SK, Audero MEP, Pontel LB, Soncini FC: **Dissecting the Salmonella response to copper**. *Microbiology (Reading)* 2007, **153**(Pt 9):2989-2997.

8.	Andersson M, Mattle D, Sitsel O, Klymchuk T, Nielsen AM, Moller LB, White SH, Nissen P, Gourdon P: **Copper-transporting P-type ATPases use a unique ion-release pathway**. *Nat Struct Mol Biol* 2014, **21**(1):43-48.

### Supplementary protocols A manual for detecting CopA of high confidence from metagenomes

#### 1. Preparation of local CopA database

1.1 The local CopA database comprises of known CopA whose function have been tested with experimental evidence or crystal structure resolved. Some were from the copper resistant strain whose genomes have been fully sequenced.
1.2 The sequences were retrieved from the Uniprot database (<https://www.uniprot.org/>). Specifically, go to the official website, search against the online database with the entry of ‘copper resistance copA’. The returned items were checked one by one, in terms of their annotation, experimental evidence and sequence features.

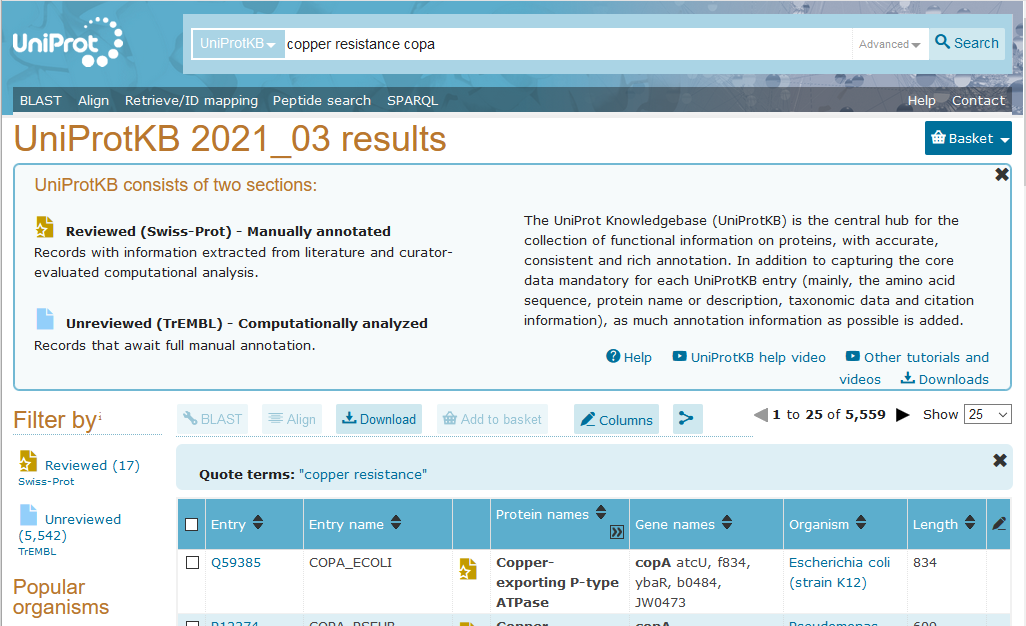

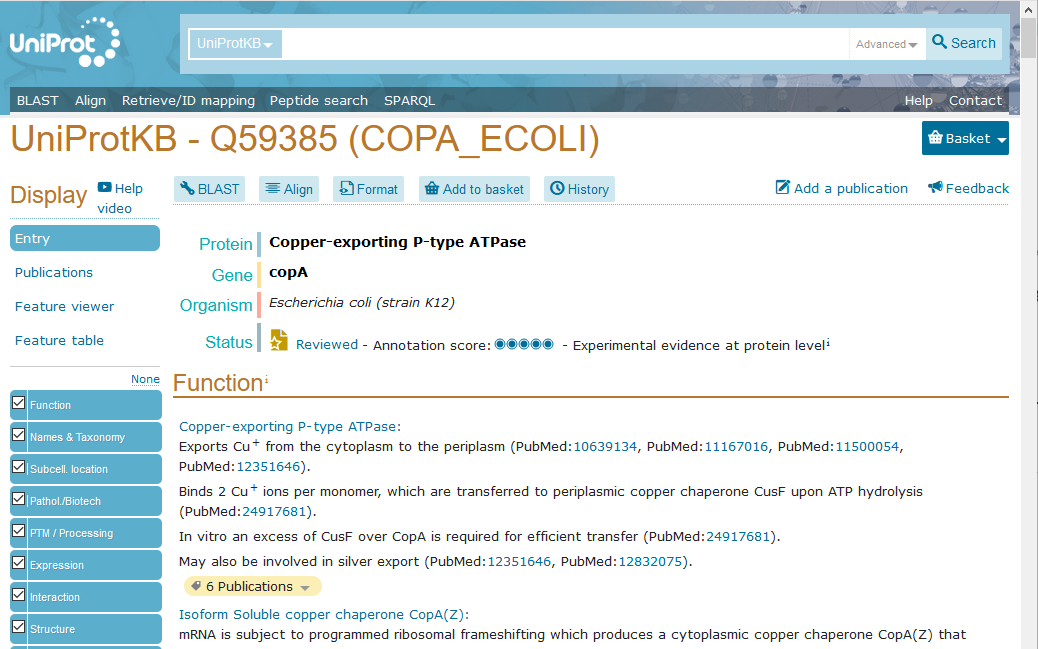

1.3 Known CopA were also searched in the literature where copper resistant bacteria were reported and their genomes were available for annotation in NCBI (https://www.ncbi.nlm.nih.gov/genome/).

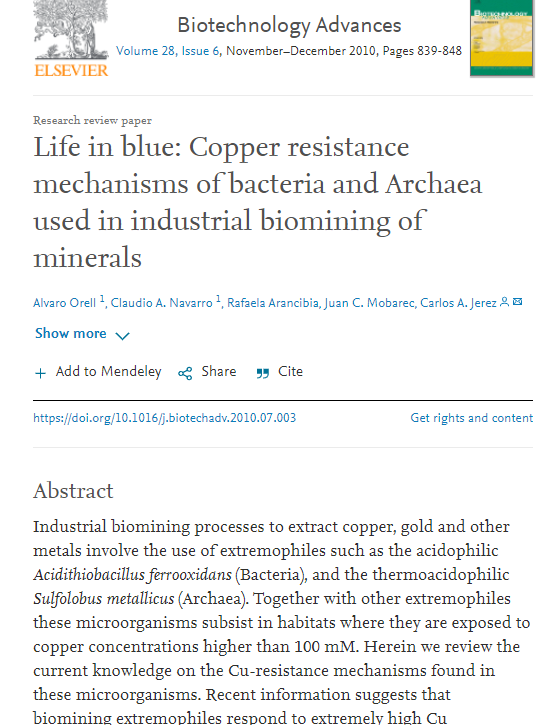

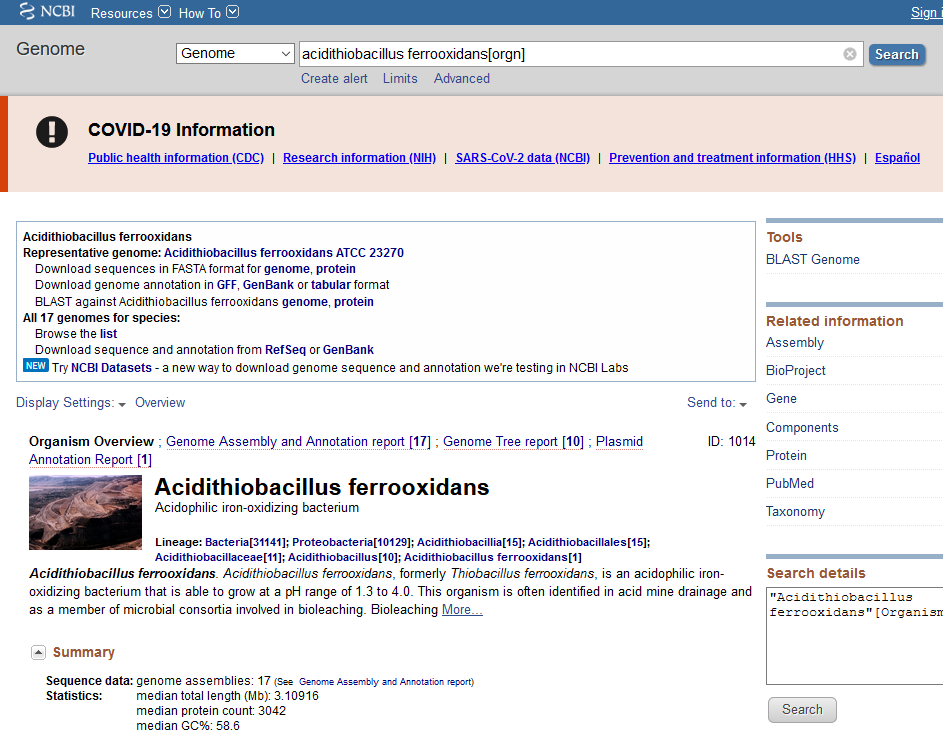

1.4 Finally a local database named ‘CopA_reviewed.fasta’ was obtained.
1.5 Phylogenetic analysis was performed to test their evolutionary relationship.

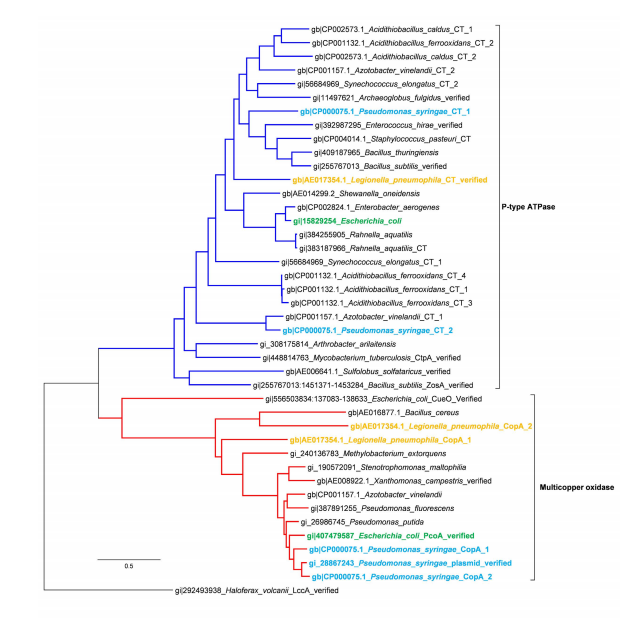

1.6 Further evolution trace analysis was done to see their length, conservative regions, key motifs for assessing novel CopA sequences.

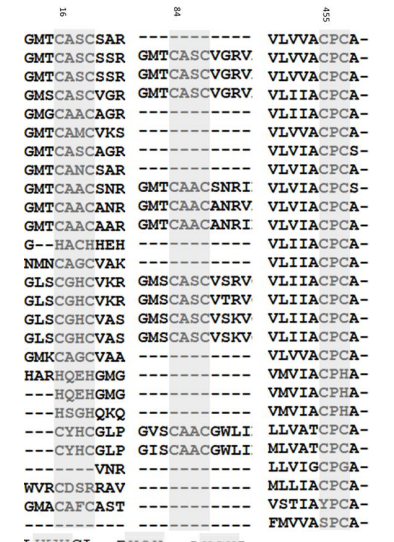

#### 2. Preparation of global metagenomes for local BLAST

2.1 Download data from MG-RAST (http://www.mg-rast.org/). Some metagenomes were downloaded as raw sequencing data, and some were processed protein assemblages.

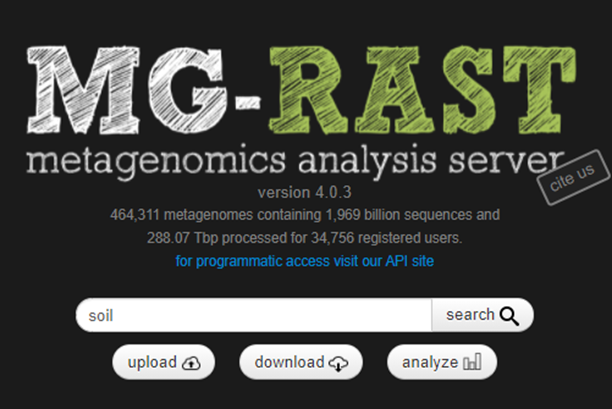

2.2 Metadata of the metagenomes including the location, data size, etc. were compiled manually for reference.

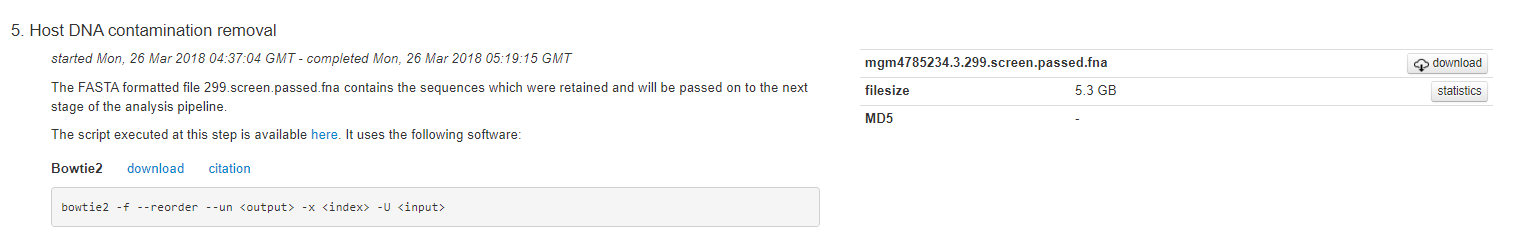

2.3 Assembly of raw data using metaWRAP.
    2.3.1 Install metaWRAP from <https://github.com/bxlab/metaWRAP> v=1.2.1
    2.3.2 Install miniconda2 by

wgethttps://repo.continuum.io/miniconda/Miniconda2-latest-Linux-x86_64.sh

bash Miniconda2-latest-Linux-x86_64.sh

2.3.3 Install metaWRAP in the conda environment

\# Note: the order is important

conda config --add channels defaults

conda config --add channels conda-forge

conda config --add channels bioconda

conda config --add channels ursky

conda install -y -c ursky metawrap-mg

\# Note: may take a while

\# To fix the CONCOCT endless warning messages in metaWRAP=1.2, run

conda install -y blas=2.5=mkl

2.3.4 Set up the conda environment

conda create -y -n metawrap-env python=2.7

conda activate metawrap-env

\# Note: the order is important

conda config --add channels defaults

conda config --add channels conda-forge

conda config --add channels bioconda

conda config --add channels ursky

conda install -y -c ursky metawrap-mg

\# Note: may take a while

\# To fix the CONCOCT endless warning messages in metaWRAP=1.2, run

conda install -y blas=2.5=mkl

2.3.5 Download the database to local

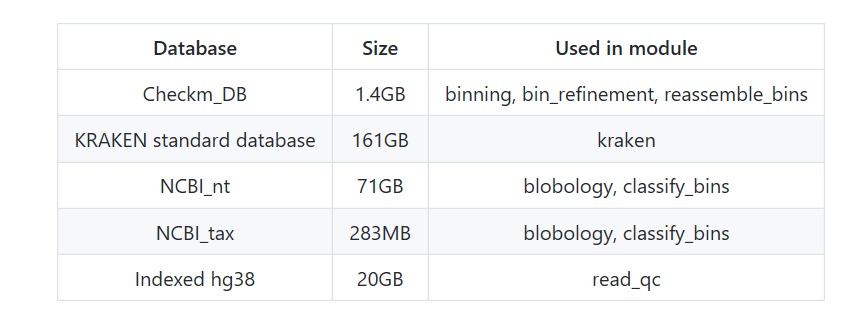

mkdir MY_CHECKM_FOLDER

checkm data setRoot

\# CheckM will prompt to to chose your storage location... Give it the path to the folder you just made.

#Now manually download the database @ https://github.com/bxlab/metaWRAP/blob/master/installation/database_installation.md
 cd MY_CHECKM_FOLDER

wget;https://data.ace.uq.edu.au/public/CheckM_databases/checkm_data_2015_01_16.tar.gz

tar -xvf *.tar.gz

rm *.gz

2.3.6 Assemble the raw data

MataWRAP -h

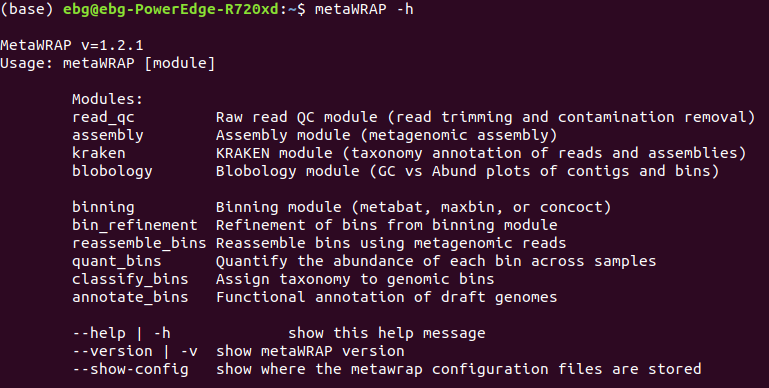

metawrap assembly -h

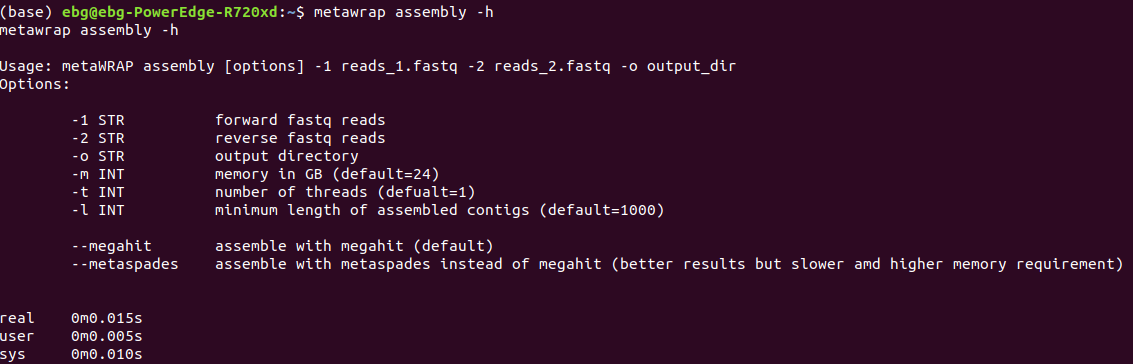

$ metawrap assembly -1 data_1.fasta -2 data_2.fasta -o output _dir

##“data.fasta” is the dataset，“data_1.fasta” is the forward sequences，“data_2.fasta” is the reverse sequences.

\# and you can view the output files here

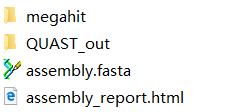

#### 3. Local BLAST for detection of potential CopA

3.1 Download software blast 2.9.0 at ftp://ftp.ncbi.nlm.nih.gov/blast/executables/blast+/LATEST/. Set up the user environmental variables. Create a new variable named BLASTDB in the user variable, and the value of the variable is C：\Blast\db. Add variable value C:\Blast\bin under the system variable “Path”.
3.2 #Format the database

makeblastdb -in CopA_reviewed.fasta -dbtype prot -title " CopA_uniprot_reviewed" -out NR

#Use NR as the local database, and assembly. fasta as subject database.

blastx.exe -db NR -query assembly.fasta -out assembly.out -evalue 0.000001 -max_target_seqs 5 -num_threads 4 -outfmt6

blastp.exe -db NR -query dirp.fasta -out dirp.out -evalue 0.000001 -max_target_seqs 5 -num_threads 4 -outfmt 6

\## A file with .out will be obtained.

3.3 Check the sequence quality

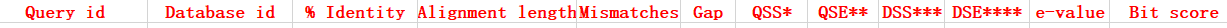

Open .out file in this excel table, filter the sequences with proper similarity and bit score.

3.4 Manually curate the sequences of high quality, and retrieve the ORFs from the sequences through online ORFfinder (https://www.ncbi.nlm.nih.gov/orffinder/) and BLAST.

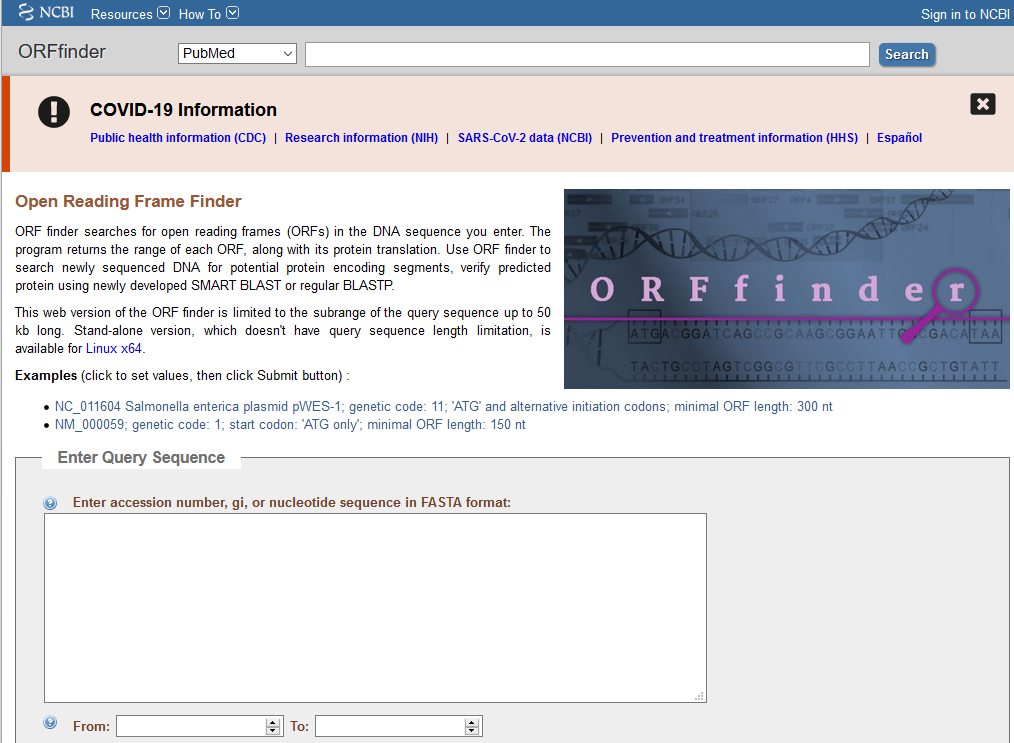

3.5 Collect the candidate *copA* genes and formed a new fasta file.

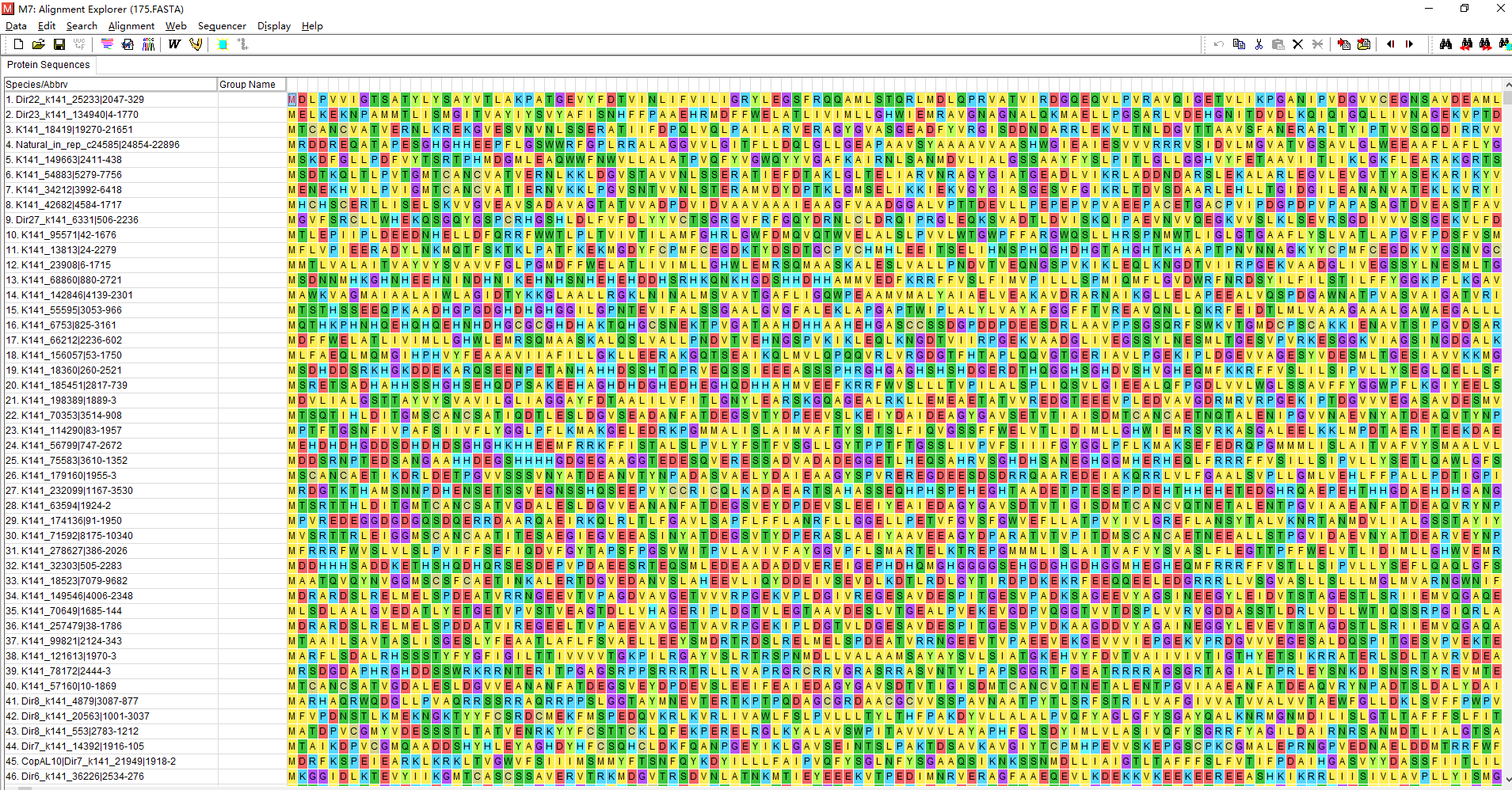

#### 4. Evolutionary trace analysis of known CopA

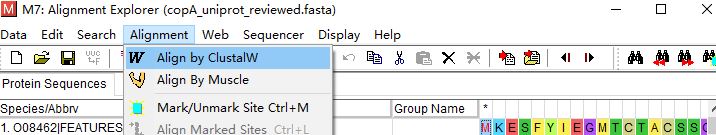

①

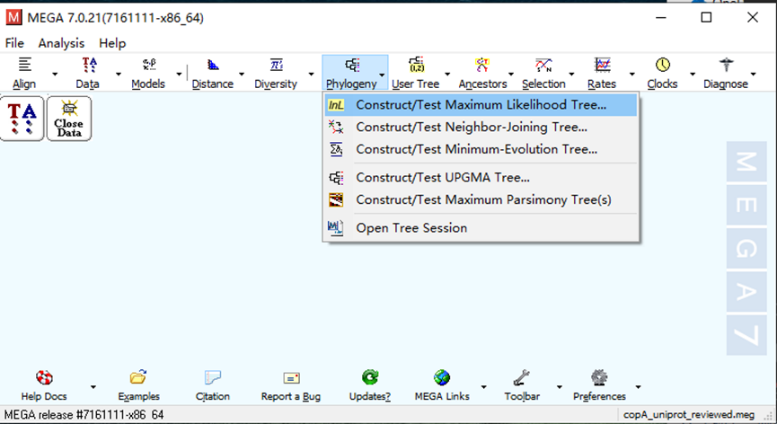

②

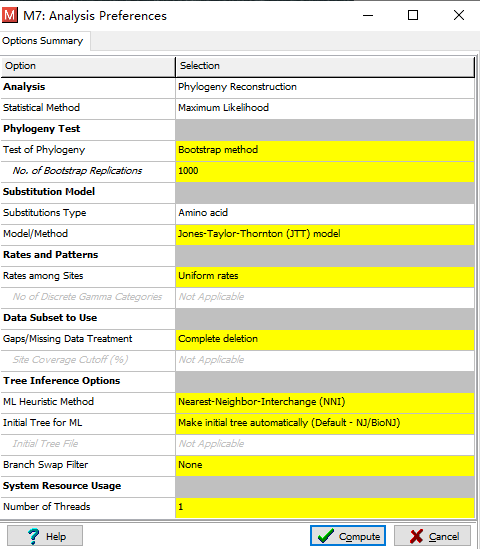

③

#### 5. Bioinformatic evaluation of potential CopA

5.1 Candidate ORFs with the length ranging from 500 to 900 amino acids were kept and subjected to the prediction of transmembrane helices using the TMHMM (http://www.cbs.dtu.dk/services/TMHMM-2.0/) online analysis platform.

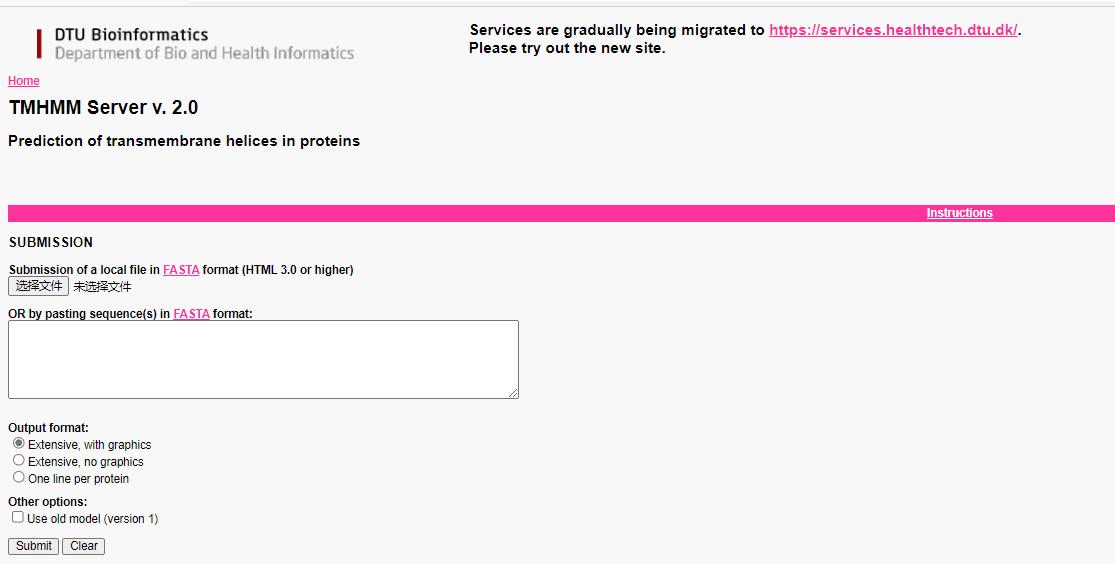

5.2 Functional domains were then predicted using Pfam (http://pfam.xfam.org/) with an E-value of ≤ 1e-6, and sequences without metal transporting domains were eliminated.

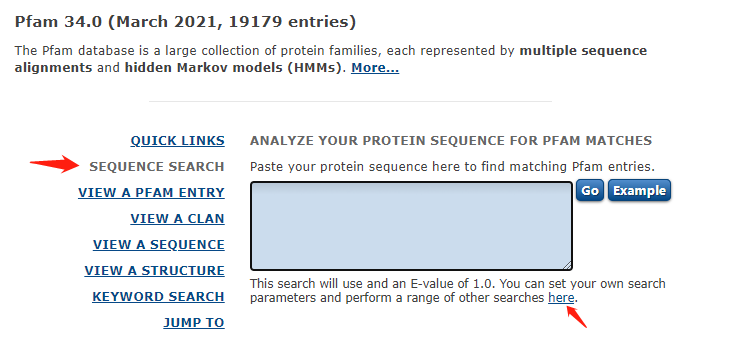

#### 6. Phylogenetic analysis of candidate CopA

Same as evolutionary trace analysis of known CopA.

#### 7. Functional genomic verification of candidate CopA (details can be found in the main text)

7.1 Chemcial sysnthesis

Gene sequence optimization in order to make sure the expression of candidate *copA*. This is the key procedure of this step.

7.2 Preparation of host, positive control and MIC test

Synthesized genes were ligated with the PTR vector (with Kna and Amp resistance) individually, and transform the vector into *copA* defective strain JW-0473-4 to verify whether the genes can restore the copper resistance of the defective strain.

7.3 Drop assay and growth test

Experimental tests on the growth of the transformed strain by growing solid LB medium with Cd supplied.

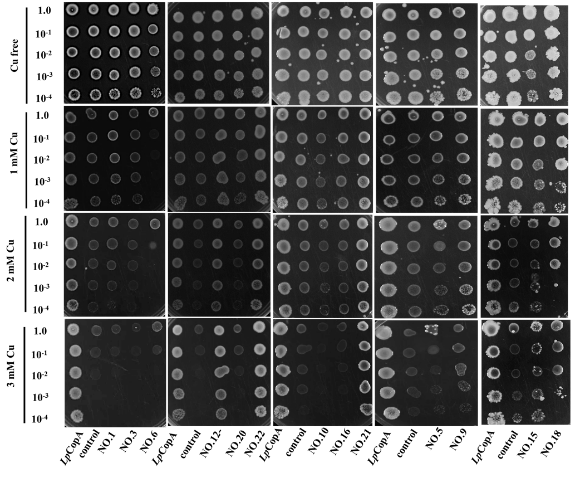

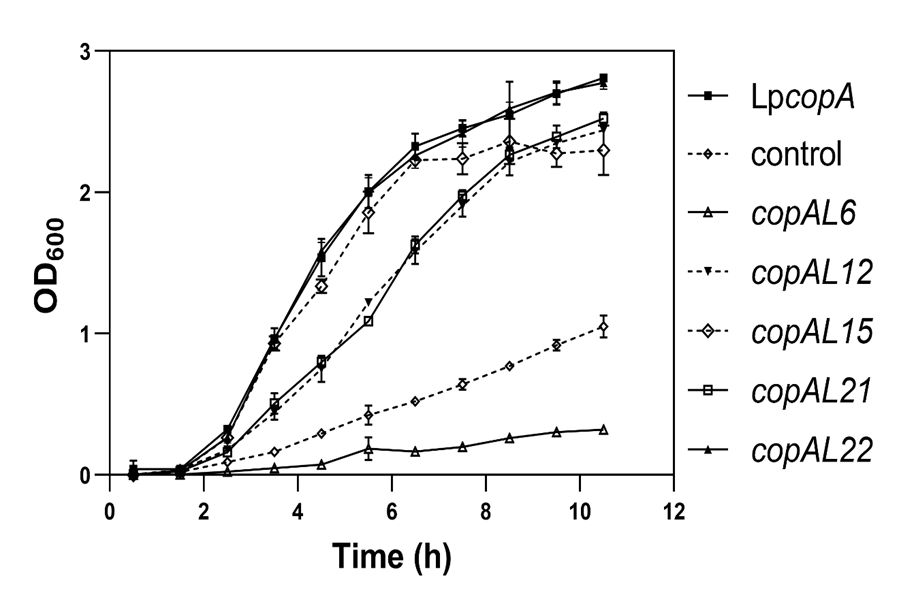
